# Pyk2 Overexpression in Postsynaptic Neurons Blocks Aβ_1-42_-induced Synaptotoxicity in a Microfluidic Co-Culture Model

**DOI:** 10.1101/2019.12.20.884205

**Authors:** Devrim Kilinc, Anaïs-Camille Vreulx, Tiago Mendes, Amandine Flaig, Diego Marques-Coelho, Maxime Verschoore, Florie Demiautte, Philippe Amouyel, Neuro-CEB Brain Bank, Fanny Eysert, Pierre Dourlen, Julien Chapuis, Marcos Romualdo Costa, Nicolas Malmanche, Frederic Checler, Jean-Charles Lambert

## Abstract

Recent meta-analyses of genome-wide association studies identified a number of genetic risk factors of Alzheimer’s disease; however, little is known about the mechanisms by which they contribute to the pathological process. As synapse loss is observed at the earliest stage of Alzheimer’s disease, deciphering the impact of Alzheimer’s risk genes on synapse formation and maintenance is of great interest. In this paper, we report a microfluidic co-culture device that physically isolates synapses from pre- and postsynaptic neurons and chronically exposes them to toxic amyloid-beta (Aβ) peptides secreted by model cell lines overexpressing wild-type or mutated (V717I) amyloid precursor protein (APP). Co-culture with cells overexpressing mutated APP exposed the synapses of primary hippocampal neurons to Aβ_1-42_ molecules at nanomolar concentrations and induced a significant decrease in synaptic connectivity, as evidenced by distance-based assignment of postsynaptic puncta to presynaptic puncta. Treating the cells with antibodies that target different forms of Aβ suggested that low molecular weight oligomers are the likely culprit. As proof of concept, we demonstrate that overexpression of protein tyrosine kinase 2 beta (Pyk2) –an Alzheimer’s disease genetic risk factor involved in synaptic plasticity and shown to decrease in Alzheimer’s disease brains at gene expression and protein levels–selectively in postsynaptic neurons is protective against Aβ_1-42_-induced synaptotoxicity. In summary, our lab-on-a-chip device provides a physiologically-relevant model of Alzheimer’s disease-related synaptotoxicity, optimal for assessing the impact of risk genes in pre- and postsynaptic compartments.

## Introduction

Alzheimer’s disease, the most common neurodegenerative disorder worldwide, is characterized by two types of brain lesions: (i) neurofibrillary degeneration due to the intracellular aggregation of abnormally hyperphosphorylated tau protein and (ii) amyloid plaques resulting from the extracellular accumulation of amyloid beta (Aβ) peptides (Nisbet *et al*., 2015). Aβ peptides are generated by the cleavage of the transmembrane amyloid precursor protein (APP) and can have different residue lengths (De Strooper, 2010). The discovery of mutations in the APP, PS1 and PS2 genes (coding for APP and presenilins 1 and 2) causing early-onset, autosomal-dominant forms of Alzheimer’s disease has profoundly influenced our understanding of the disease, and has placed Aβ peptides at the center of the pathophysiological process. According to the “amyloid cascade hypothesis”, the accumulation of Aβ peptides is the triggering toxic condition that induces the development of neurofibrillary degeneration and thus neuronal death (Hardy and Selkoe, 2002).

Aβ_1-42_ species have been the principal focus of research (Stine *et al*., 2003) since Aβ_1-42_ is more prone to oligomerize (Dahlgren *et al*., 2002; Resende *et al*., 2008) and oligomers of Aβ_1-42_ are more toxic than its monomeric or fibrillary forms, and other Aβ species (Deshpande *et al*., 2006; Ferreira *et al*., 2012; Spires-Jones and Hyman, 2014). Oligomeric Aβ_1-42_ is in a dynamic equilibrium with the monomeric forms and fibrils, and has been proposed to be the main promoter of amyloid plaques (Benilova *et al*., 2012). Although the central role of the Aβ peptide burden as initially enunciated in this hypothesis is strongly debated, several lines of evidence indicate that Aβ peptides are still a key actor of the disease at least *via* their oligomeric forms. In particular, the Aβ oligomer toxicity has been linked with synapse dysregulation and loss (Brody and Strittmatter, 2018).

Synapse loss is a major pathological correlate of cognitive deficits in Alzheimer’s disease (Lansbury, 1999) and is observed at the earliest stage of the disease (Scheff *et al*., 2007). Several mechanisms have been proposed to explain Aβ-induced synaptotoxicity: (i) membrane-disrupting activity at high concentrations (Sepulveda *et al*., 2010); (ii) deleterious pruning of synapses by microglia activation (Hong *et al*., 2016); (iii) direct interaction of oligomeric forms with postsynaptic receptors, such as ionotrophic or metabotropic glutamate receptors (Wang *et al*., 2004; Um *et al*., 2012). In addition, the new genetic landscape of Alzheimer’s disease, resulting from the advent of the genome wide association studies (GWAS) (Lambert *et al*., 2013; Kunkle *et al*., 2019), highlights synaptic (dys)regulation: several among the dozens of genes/loci identified to be associated with Alzheimer’s disease risk, *e.g., BIN1, CD2AP, FERMT2* and *PTK2B*, have been shown to modulate synaptic functions in the physio- and/or pathophysiological contexts (Giralt *et al*., 2017; Eysert *et al*., 2019; Ojelade *et al*., 2019; Salazar *et al*., 2019; Schurmann *et al*., 2019). As a result, we recently proposed a genetically-driven synaptic failure hypothesis, based on the genetic and post-GWAS data (Dourlen *et al*., 2019). In this context, Aβ toxicity is one of the elements involved in synapse failure and it may be driven by specific Alzheimer’s disease genetic risk factors.

However, assessing such an hypothesis requires a number of considerations to be taken into account: (i) most *in vitro* models of Aβ toxicity use synthetic Aβ oligomers at non-physiological concentrations, even though synthetic fibrils are structurally different from Aβ fibrils obtained from Alzheimer’s brains (Kollmer *et al*., 2019); (ii) only a few of the GWAS-defined genes have been analyzed in a (physiological and/or pathophysiological) synaptic context; (iii) GWAS-defined genes may have different effects when expressed in the pre- or postsynaptic neurons. For example, protein tyrosine kinase 2 (Pyk2), product of Alzheimer’s disease risk gene *PTK2B*, directly interacts with postsynaptic scaffold proteins (Bartos *et al*., 2010), regulates dendritic spine morphology (Giralt *et al*., 2017), and is involved in synaptic plasticity through regulating postsynaptic NMDA receptors *via* activation of Src (Huang *et al*., 2001). Recent studies based on Alzheimer’s disease mouse models linked Pyk2’s effects to amyloid pathology, but reported contradictory data on whether lack of Pyk2 was detrimental or protective (Giralt *et al*., 2018; Salazar *et al*., 2019).

With this background, we have developed a microfluidic co-culture device, based on existing tricompartment designs that physically isolate synapses and provide exclusive access to pre- and postsynaptic neurons (Taylor *et al*., 2010; Virlogeux *et al*., 2018). This device permits not only the induction of synaptotoxicity *via* physiologically-relevant concentrations of Aβ molecules secreted by cells stably overexpressing human APP, but also the analysis of synaptic density as a function of over- or underexpression of genetic risk factors in pre- and/or postsynaptic neurons. In this paper, we characterized Aβ-induced synaptotoxicity in primary neurons upon co-culture with cells overexpressing mutated APP and assessed the impact of Pky2 overexpression in postsynaptic neurons, as a proof of concept.

## Experimental

### Oligimerization of synthetic Aβ peptides

Aβ peptides were oligomerized according to established protocols (Stine *et al*., 2003; Chang *et al*., 2012) with minor modifications. The inactive control peptide (Aβ_42-1_; Abcam; Cambridge, UK) has the same composition as the Aβ_1-42_ peptide (California Peptide Research; Napa, CA) but with an inverted amino-acid sequence, and has been widely used as control for oligomeric Aβ_1-42_ since it is also prone to oligomerization (Walsh *et al*., 2002; Xiong *et al*., 2007; Mairet-Coello *et al*., 2013). Aβ_1-42_ and Aβ_42-1_ were treated with hexafluoroisopropanol (Sigma-Aldrich, Saint Louis, MO) to maintain the oligomeric structure and to reduce fibril formation (Stine *et al*., 2003), according to manufacturer’s instructions. The peptides were resuspended in 1 ml hexafluoroisopropanol and incubated for 1 h at RT, with occasional moderate vortexing, followed by sonication in water bath (Branson; Emerson Electric; St. Louis, MO) for 10 min. The solution was aliquoted into microcentrifuge tubes, let to evaporate in a chemical fume hood, dried with SpeedVac system (Thermo Fisher Scientific, Waltham, MA) for 30 min, and stored as desiccated peptide at −80°C. To produce oligomers, lyophilized, hexafluoroisopropanol-treated aliquots of both peptides were resuspended in DMSO to reach 5 mM, mixed by pipetting, sonicated in water bath for 10 min, diluted to 100 μM in ice-cold D-PBS, followed by 30 s vortexing and 1 h incubation at 25°C. The solutions were aliquoted into microcentrifuge tubes and stored at −20°C for up to four weeks. Before adding to neurons, oligomeric peptide stocks were thawed, serial diluted to 100 nM in 2 % DMSO in PBS. Aβ_1-42_ concentrations up to 1 μM are considered non-lethal (Kelly and Ferreira, 2007; Kuperstein *et al*., 2010).

### Microfluidic device design and fabrication

The microfluidic co-culture device was designed based on a previous tricompartmental neuron culture device (Kilinc *et al*., 2014). The device consists of a 300 μm wide, 7.4 mm long central channel flanked by two 750 μm wide, 3.6 mm long side channels. All three channels are *ca*. 100 μm high. The left side channel (termed presynaptic) and the central channel (termed synaptic) are interconnected *via* 4 μm high, 450 μm long parallel microchannels that narrow from an entry width of 10 μm to an exit width of 3 μm. The right side channel (termed postsynaptic) and the synaptic chamber are also interconnected *via* parallel microchannels with identical dimensions, except that they were 75 μm long. One end of the synaptic chamber bifurcates into two branches, one of which terminates in a triangular shape. This terminus is connected to a diamond-shaped co-culture chamber (based on a previous design) (Kilinc *et al*., 2016) *via* 4 μm high, 10 μm wide, and 100 μm long parallel microchannels.

Master patterns were fabricated at the IEMN (Lille, France) *via* two-step photolithography (Blasiak *et al*., 2015). *Ca*. 4 mm high polydimethysiloxane (PDMS) pads were replica molded. Access wells were punched at the termini of the central channel and the co-culture chamber, and of the side channels using 3 mm and 4 mm biopsy punches (Harris Unicore), respectively. The devices were permanently bonded to 24 mm × 50 mm glass coverslips (Menzel) *via* O_2_ plasma (Diener, Ebhausen, Germany). Prior to cell culture, the devices were sterilized under UV light (Light Progress, Anghiari, Italy) for 30min, treated with 0.1 mg/mL PLL overnight, and rinsed with PBS.

### Primary neuron culture

Culture media and supplements were from Thermo Fisher, unless mentioned otherwise. Primary neurons were obtained from P0 rats, according to previously described procedures (Sartori *et al*., 2019). Briefly, cortices and hippocampi were isolated from new-born rats, washed with ice-cold dissection medium (HBSS supplemented with HEPES, sodium pyruvate, and penicillin/streptomycin), and trypsinized (2.5%; at 37°C for 10 min). Trypsin was inactivated with dissociation medium –MEM supplemented with fetal bovine serum (FBS), Glutamax, D-glucose (Sigma), MEM vitamins, and Pen/Strep–followed by DNase (5 mg/ml; Sigma) incubation for 1 min and wash with dissection medium. Media was replaced by dissociation medium and tissue was triturated with a fire-polished cotton-plugged Pasteur pipette to obtain a homogenous cell suspension, followed by centrifugation (200× *g* for 5 min). Cells were resuspended in culture medium (Neurobasal A supplemented with Glutamax and B27 neural supplement with antioxidants), counted, and plated at a density of 100,000 cells/cm^2^ in 6- and 24-well plates for immunoblots and in 10 cm Petri dishes for synaptosome extraction. Plates were pre-coated with 0.1 mg/mL poly-l-lysine (Sigma) in 0.1M borate buffer (0.31% boric acid, 0.475% sodium tetraborate, pH = 8.5; Sigma) overnight at 37°C and rinsed thoroughly with water. Alternatively, cells were plated in pre-coated 384-well plates at 50,000 cells/cm^2^ (*ca*. 4000 cells/well) and in microfluidic devices at a density of *ca*. 8×10^5^ cells/cm^2^. After 20-24 h, culture media was replaced with supplemented Neurobasal A medium. 0.1% ethylenediaminetetraacetic acid (EDTA; in H2O) was added to the Petri dishes containing microfluidic devices to minimize evaporation. Cultures were maintained in a tissue culture incubator (Panasonic; Osaka, Japan) at 37°C and 5% CO_2_.

### Immunoblotting

Neurons were harvested in minimum volume of 200 μl/well in 6-well plates, in ice-cold lysis buffer as described earlier (Chapuis *et al*., 2017). Lysates were mixed with 4× LDS sample buffer (Novex; Thermo Fisher) and 10× reducing agent (Novex), sonicated, and boiled at 95°C for 10 min. Samples were loaded at maximum volume into 1.5 mm, 10-well, 4-12% Bis-Tris pre-cast NuPage gels (Novex), along with 5 μl MW marker (Novex Sharp pre-stained protein standard, Thermo Fisher). The gel was run with 2-(N-morpholino)ethanesulfonic acid (MES) running buffer at 200 V for 45 min and transferred to 0.22 μm nitrocellulose membranes using the Trans-Blot Turbo transfer system (BioRad, Hercules, CA) using mixed MW method at 1.3 A and 25 V for 7 min. Membranes were blocked in TNT (0.05% Tween 20, 20 mM Tris-Base, 150 mM NaCl, pH = 8.0) containing 5% non-fat dry milk for 1 h at RT and washed 3× in TNT. Membranes were incubated with the following primary antibodies in SuperBlock T20 blocking buffer (Thermo Scientific) at 4°C overnight and washed 3× in TNT: rabbit anti-phospho-PTK2B (Cell Signaling Technology 3291; 1/1000; Danvers, MA), rabbit anti-PTK2B (Sigma P3902; 1/1000), and mouse anti-β-actin (Sigma A1978; 1/5000). Membranes were then incubated with horseradish peroxidase (HRP)-conjugated secondary antibodies (HRP-anti-Mouse and HRP-anti-rabbit; 1:5000; Jackson ImmunoResearch, West Grove, PA) in TNT containing 5% non-fat dry milk for 1 h at RT and washed 3× in TNT. The membrane was revealed through chemiluminescence (Luminata Classico, EMD Merck Millipore) and imaged with Amersham Imager 600 (GE Healthcare, Mississauga, Canada). The images were quantified with ImageQuantTL Software (GE Healthcare).

### Culture of CHO cells and analysis of their media

CHO cell lines (CHO-pcDNA4, -APP^WT^ and -APP^LDN^) were maintained according to previously described procedures (Guillot-Sestier *et al*., 2012). Cells were grown in CHO growth medium: DMEM/Ham’s F-12 1:1 medium, supplemented with 10% heat-inactivated FBS, 0.2% Pen/Strep, 2% HT supplement, and 300 μM Proline (Sigma). To stimulate Aβ production, the growth medium was replaced with CHO-NBA medium: phenol red-free Neurobasal A (Gibco) supplemented with 0.2% Pen/Strep, 2% HT supplement, and 300 μM Proline.

For media collection, cells were grown in 10 cm Petri dishes or in 6-well plates until they reached 80% confluence, at which point the maintenance medium was rinsed with PBS and replaced with the stimulation medium. After 48 h of stimulation, the medium was collected into 15 mL Falcon tubes and centrifuged at 4000× *g* and 4°C for 10 min to remove the debris. The supernatant was loaded into a 3 kDa spin column (Amicon Ultra; Merck), equilibrated with Neurobasal (without serum or Phenol Red) at 4000× *g* and 4°C for 10 min, and concentrated at 4000× *g* and 4°C for 1 h. Western blotting of conditioned media was performed as described with the following exceptions: the transferred membrane was boiled for 5 min in PBS and Luminata Crescendo (Millipore) was used as the HRP substrate. Anti-Aβ_1-42_ (clone 6E10; 1:1000; Sigma) was used as primary antibody.

### Exposure of neurons to conditioned media

Total protein concentration in the conditioned media collected from different CHO cell lines (CHO-pcDNA4, -APP^WT^ and - APP^LDN^) was assessed using the Pierce BCA Protein Assay Kit (Thermo Fisher) and adjusted to 100 μg/μL with supplemented CHO-NBA medium. 10 μL of conditioned media (per well) was added to primary neuronal cultures grown in 384-well plates (containing 40 μL of culture medium per well) at DIV14, DIV18, DIV19, DIV20, and DIV21 (6 h prior to media collection). Media from these wells (eight wells per condition) were collected into a new 384-well plate prior to quantification of Aβ peptides.

### Co-culture of neurons with CHO cells

CHO cells were seeded in the co-culture chamber at a density of *ca*. 1.3×10^5^ cells/cm^2^ and maintained in a tissue culture incubator (5% CO_2_; 37°C) in CHO medium. On the day of primary neuron culture, the medium was replaced with CHO-NBA medium supplemented with 1% Glutamax and 2% B27 neural supplement with antioxidants. Primary neurons were seeded in the pre- and postsynaptic chambers. At DIV1, the medium in all wells was replaced with fresh supplemented CHO-NBA medium. Every 3-4 days, the medium in the access wells of the co-culture chamber was replaced with fresh medium, whereas the medium in other wells were only topped up with fresh medium.

### Lentiviral transductions

At DIV7, neurons cultured in the postsynaptic chamber were transducted with lentiviruses according to established methods.(Sartori *et al*., 2019) To avoid transduction of CHO cells and the neurons in the presynaptic chamber, a hydrostatic pressure gradient was formed across the microchannels separating synaptic and postsynaptic chambers. The following lentiviruses were used for transduction: Mission shRNA vectors (Sigma) pLenti6/Ubc/v5-DEST (Invitrogen, Carlsbad, CA) empty (Mock) or including human PTK2B cDNA sequences, synthesized *via* the GeneArt service (Thermo Fisher). LifeAct-Ruby (pLenti.PGK.LifeAct-Ruby.W; RRID: Addgene_51009) and LifeAct-GFP (pLenti.PGK.LifeAct-GFP.W; RRID: Addgene_51010) were kind gifts from Rusty Lansford. Viral transduction was performed at multiplicity of infection (MOI) of four. Constructs were diluted in pre-warmed, supplemented CHO-NBA medium containing 2 μg/mL (5×) Polybrene (hexadimethrine bromide; Sigma). Media from pre- and postsynaptic wells were collected in a common tube. 25 μL, 15 μL, and 20 μL of the collected media were returned to each presynaptic, synaptic, and postsynaptic well, respectively. 10 μL of virus suspension was added to one of the postsynaptic wells. Neurons were incubated with viral particles for 6 h before the wells were topped up with the remainder of the collected media. Co-cultures were maintained as described earlier.

### Alpha-LISA measurements

Alpha-LISA is a highly sensitive, quantitative assay based on biotinylated antibody bound to streptavidin-coated donor beads and antibody-conjugated acceptor beads. In the presence of the analyte, the beads come into close proximity such that the excitation of the donor beads triggers a cascade of energy transfer in the acceptor beads, resulting in a sharp peak of light emission at 615 nm. We used Alpha-LISA kits specific to human Aβ_1-X_ and Aβ_1-42_ (AL288C and AL276C, respectively; PerkinElmer, Waltham, MA) to measure the amount of Aβ_1-X_ and Aβ_1-42_ in culture media. The human Aβ analyte standard was diluted in CHO-NBA medium. For the assay, we first added 5 μL of cell culture supernatant or standard solution into an Optiplate-96 microplate (PerkinElmer). We then added 5 μL of 10× mixture including acceptor beads and biotinylated antibody. Following incubation at RT for 60 min, we added 40 μL of 1.25 × donor beads and incubated at RT for 60 min. We measured the fluorescence using an EnVision-Alpha Reader (PerkinElmer) at 680 nm excitation and 615 nm emission wavelengths. In experiments where conditioned media was added to primary neurons in 384-well plates, 2 μL of collected media was transferred to an Optiplate-384 (PerkinElmer) and Alpha-LISA was performed using reduced volumes: 2 μL of sample or standard, 2 μL of acceptor beads and biotinylated antibody mix, and 16 μL of donor beads.

### ELISA measurements

The sandwich ELISA was performed according to the manufacturer’s protocol. We used microtiter plates pre-coated with anti-human Aβ_35-40_ antibody (clone 1A10; RE59781, IBL) and anti-human Aβ_38-42_ (clone 1C3; RE59721, IBL) to detect Aβ1-40 and AβX-42, respectively. Plates were incubated overnight at 4°C with 100 μL cell culture supernatant or with standards. The bound antigen was detected by incubating the wells with 100 μL of 30× mixture containing the HRP-conjugated anti-human Aβ antibodies (clone 82E1 for Aβ_1-40_ and clone 12B2 for Aβ_X-42_) for 60 min at 4°C. Signal was revealed by incubating with the TMB substrate for 30 min at RT in the dark and stopping the enzymatic reaction with the TMB stop solution containing 1N H_2_SO_4_. Signal intensity was read immediately at 405 nm *via* a microplate reader (PowerWave XS2, BioTek Instruments, Winooski, VT).

### Neuronal viability assay using conventional co-cultures

Cell culture inserts (Falcon 353104; Corning; Corning, NY) were placed in 24-well plates and pre-incubated with 500 μl CHO growth medium for 30 min. CHO cells (CHO-pcDNA4, CHO-APP^WT^, and CHO-APP^LDN^) were plated at a density of 4,000 cells/insert and incubated at 37°C and 5% CO_2_ for 3 d, during which the cells reached confluence. CHO growth medium was then replaced with CHO-NBA medium supplemented with 1% Glutamax and 2% B27 with antioxidants and primary neurons were plated in the 24-well plate containing the inserts at a density of 2×10^5^ cells/well. As control conditions, primary neurons were plated in the wells with inserts without CHO cells and CHO cells were plated in inserts without primary neurons, under otherwise identical conditions. After 24 h, the culture media was replaced with fresh, supplemented CHO-NBA medium. Co-cultures were maintained in a tissue culture incubator at 37°C and 5% CO_2_ for 14 d. 200 μl medium was collected from the wells at DIV1, DIV7, and DIV14 for lactate dehydrogenase (LDH) release assay. LDH Cytotoxicity Detection Kit (Takara Bio; Saint-Germain-en-Laye, France) was used according to the manufacturer’s protocol. Briefly, collected culture media was incubated with reaction mixture at 1:1 ratio for 30 min (in dark, at RT) and the LDH enzymatic activity was subsequently measured with a microplate reader at 490 nm (PowerWave XS2; BioTek Instruments). Cell viability was calculated as a percentage of the difference between negative controls (primary neurons without CHO cell co-culture treated with 0.1% Triton X-100 for 5 min) and positive controls (primary neurons without CHO cell co-culture). The effect of CHO cell co-culture on neuronal viability was determined by subtracting the respective CHO cell mono-culture signal from the co-culture signal. The presence of Aβ_1-42_ in the conventional co-culture media was measured via alpha-LISA as described.

### Immunocytochemistry and microscopy

Co-cultures were fixed at DIV14 in PBS containing 4% paraformaldehyde for 20 min at RT and permeabilized with 0.3% (v/v) Triton X-100 in PBS for 5 min. After blocking in 5% (w/v) normal donkey serum, samples were incubated overnight at 4°C with the following antibodies: mouse anti-MAP2 (188011; Synaptic Systems, Göttingen, Germany); chicken anti-Homer (160006; Synaptic Systems) and rabbit anti-PYK2 phospho-Y402 (ab4800; Abcam, Cambridge, UK). Cells were rinsed with PBS and incubated with the following secondary antibodies for 2 h at RT: Dylight 405 Donkey anti-mouse (715-475-151; Jackson); AlexaFluor 647 donkey anti-rabbit (711-605-152; Jackson) and AlexaFluor 594 Donkey anti-chicken (703-585-155; Jackson). Cells were rinsed with PBS and incubated with mouse monoclonal anti-Synaptophysin 1 pre-labeled with Sulfo-Cyanine 2 (101011C2; Synaptic Systems) for 2 h at RT. Cells were rinsed with PBS and microfluidic devices were topped with 90% glycerol.

Samples were imaged with a LSM 880 confocal microscope (Zeiss, Oberkochen, Germany) equipped with a 63× 1.4 NA objective. Images were acquired at zoom 2 in *z*-stacks of 0.5 μm interval. Typically, four images were acquired per device from the synapse chamber near the postsynaptic chamber such the image contains multiple dendrites. Images were deconvoluted using AutoQuantX3 software (BitPlane, Zurich, Switzerland) for synaptic connectivity analysis.

In a subset of experiments, neurons were plated only in the presynaptic chamber, only in the postsynaptic chamber, or only in the synaptic chamber. Neurons were fixed and immunostained at DIV14 as described to reveal nuclei, GFAP (ABD95; Millipore) and β3-tubulin (MAB1637, Millipore). All chambers were imaged using a Zeiss AxioObserver Z1 epifluorescense microscope equipped with a Prime 95B Scientific CMOS (Photometrics, Tucson, AZ) camera and 32× objective. The integrated density (pixel values × area) of the β3-tubulin signal was quantified in all chambers, at a distance between 50 and 200 μm from the entries and exits of the microchannels. Neurite penetration rates for short and long microchannels (in the forward and reverse directions) were estimated by taking the ratio of the β3 -tubulin integrated density signals obtained from the emitting and receiving chambers. The ratio between the penetration rates between forward and reverse directions was defined as the directionality ratio.

### Quantification of synapse integrity

We developed an image analysis workflow based on image segmentation using Imaris software (BitPlane) and assignment of postsynaptic signals to the nearest presynaptic signal using a custom Matlab (Mathworks; Natick, MA) code. Briefly, signals obtained for pre- and postsynaptic structures were first deconvoluted in Autoquant X3 (Media Cybernetics; Rockville, MD) and segmented into distinct volumes using the surfaces function of Imaris in the batch mode that permits the same parameters to be applied to all images (Supplementary Fig. 1A-D). Postsynaptic spots were then assigned to the nearest presynaptic spot (according to the 3-D Euclidean distance between intensity centers) within a predefined cut-off distance (Supplementary Fig. 1E). It is important to note that pre- and postsynaptic structures are smaller than the axial resolution of the confocal microscope; the distances measured are thus not true physical distances between synaptic structures, but the distances between respective intensity centers. We empirically determined the cut-off distance by testing a range of values on a large set of control cultures (Supplementary Fig. 1F-G). The fraction of Syp puncta with at least one Homer assignments and the average number of Homer assignments per Syp were determined to be the most robust read-outs of synapse connectivity. The numbers of assigned pre- and postsynaptic puncta per image area (in the *xy*-plane) were also provided. Note that a small fraction of control samples for CHO-APP^WT^ and CHO-APP^LDN^ co-cultures (three and two devices, respectively) do overlap in the data reported.

### Analysis of phospo-Pyk2 signals relative to synapses

We extended the image analysis workflow to analyze the positions of phospho-Pyk2 (Tyr402) (p-Pyk2) puncta relative to the positions of identified synapses. Postsynaptic spots were first assigned to the nearest presynaptic spot as described. For each postsynaptic-to-presynaptic assignment, or “synapse”, we defined the midpoint as being equidistant to pre- and postsynaptic puncta. We next defined pre- and postsynaptic zones, by spanning two right circular cones with 45° polar angle where the midpoint is the apex. We then associated p-Pyk2 puncta with the nearest “synapse” as long as it was within a pre-defined cut-off distance from the midpoint and categorized these associations as presynaptic, postsynaptic, or other (neither pre-nor postsynaptic). We then calculated the average number of p-Pyk2 puncta associated with each “synapse”. As each “synapse” inherently consists of a Syp and Homer pair, a paired statistical test was justified for this analysis.

### Synaptosome extraction

To verify the presence of proteins at the synaptic level we conducted subcellular fractionation as previously described (Frandemiche *et al*., 2014). Briefly, cortical neurons were cultured in 10 cm Petri dishes as described (3.5-4.0 × 10^7^ neurons per condition). At DIV13, neurons were exposed to CHO-APP^WT^ or CHO-APP^LDN^ media for 18 h (final Aβ_1-X_ concentration: 40 nM). At the end of this treatment, neurons were lysed, reconstituted in a solution (0.32 M sucrose and 10 mM HEPES, pH = 7.4), and centrifuged at 1000× *g* for 10 min to remove nuclei and debris. The supernatant was centrifuged at 12,000× *g* for 20 min to remove the cytosolic fraction. The pellet was reconstituted in a second solution (4 mM HEPES, 1 mM EDTA, pH = 7.4) and was centrifuged 2× at 12,000× *g* for 20 min. The new pellet was reconstituted in a third solution (20 mM HEPES, 100 mM NaCl, 0.5% Triton X-100, pH = 7.2) for 1 h at 4°C and centrifuged at 12,000× *g* for 20 min. The supernatant collected corresponds to the non-postsynaptic density (PSD) fraction (Triton-soluble). The remaining pellet was reconstituted in a fourth solution (20 mM HEPES, 0.15 mM NaCl, 1% Triton X-100, 1% deoxycholicacid, 1% SDS, pH = 7.5) for 1 h at 4°C and was centrifuged at 10,000× *g* for 15 min to obtain a supernatant containing the PSD fraction (Triton-insoluble). The fractions obtained were then analyzed by WB. There was no difference in the Syp signal in the non-PSD fraction and the PSD95 signal in the PSD fraction for neurons treated with CHO-APP^WT^ and CHO-APP^LDN^ media, suggesting that pre- and postsynaptic proteins were not affected by this treatment paradigm (Supplementary Fig. 2).

### Analysis of gene expression alterations in human brain samples

RNA sequencing (RNAseq) data from the Mount Sinai/JJ Peters VA Medical Center Brain Bank (MSBB) (Wang *et al*., 2018), the ROSMAP database (De Jager *et al*., 2018), and the Mayo Clinic whole genome and transcriptome data (Allen *et al*., 2016) were downloaded from AMP-AD Knowledge Portal (https://adknowledgeportal.synapse.org/) according to the terms and conditions concerning the use of the data. Data were aligned using pseudoaligner Kallisto version 0.43.1 (Bray *et al*., 2016) using a pre-built index to align fastq files. Differential gene expression analysis was performed using DESeq2 (Love *et al*., 2014). First, a DESeq2 object was created using *DESeqDataSetFromTximport* function and rows with sum of all counts less than 10 were filtered out. Next, *DESeq* function was used with default parameters. Temporal cortex was analyzed in the Mayo Clinic dataset (82 cases and 78 healthy controls). Dorsolateral prefrontal cortex was analyzed in the ROSMAP dataset (222 cases and 201 controls). The following brain areas were analyzed in the MSBB dataset: Brodmann area (BA) 10, which corresponds to the anterior prefrontal cortex (105 cases and 71 controls); BA 22, which is part of the Wernicke’s area in the superior temporal gyrus (98 cases and 61 controls); BA 36, which corresponds to the lateral perirhinal cortex (88 cases and 64 controls); and BA 44, which corresponds to the inferior frontal gyrus (90 cases and 63 controls).

### Analysis of phosphorylated and total Pyk2 levels in human brain samples

The brain samples were collected through a brain donation program dedicated to neurodegenerative dementias coordinated by the NeuroCEB Brain Bank Network. The informed consent for post-mortem examination and research studies was signed by the legal representative of each patient in patient’s name, as allowed by the French law and approved by the local ethics committee and the brain bank has been officially authorized to provide samples to scientists (agreement AC-2013-1887). All procedures performed in this study involving human participants were in accordance with the ethical standards of the institutional research committees and with the 1964 Helsinki declaration. The brain banks fulfill criteria from the French Law on biological resources including informed consent, ethics review committee and data protection (article L1243-4 du Code de la Santé publique, August 2007). The Neuro-CEB brain bank (BioResource Research Impact Factor number BB-0033-00011) has been declared to the Ministry of Research and Higher Education, as required by French law.

Assessment of Alzheimer’s disease-related neurofibrillary pathology (Braak stage) was performed according to published procedures (Thierry *et al*., 2020) by analyzing post-mortem brain tissue samples of 28 individuals (Table S1) *via* immunohistochemistry against Aβ deposits (Dako M0872; clone 6F/3D; Agilent, Santa Clara, CA) and against hyperphosphorylated Tau at Ser202/Thr205 (clone AT8; Thermo Fisher) (Braak *et al*., 2006). Lysis buffer containing, trizma-base 20 mM; NaCl 150 mM; cOmplete Protease Inhibitor Cocktail 1X and 1% Triton X-100, was added to single pieces of whole brain tissue (~100 mg) at a ratio of 5 μL per 1 mg of tissue. Brain samples were homogenized by beads beating using a precellys soft tissue CK14 2 mL (3× 30 s at 6500 rpm). The lysate was then centrifuged at 4000 rpm for 15 min at 4°C. 50 μL from the supernatant was used for analysis. Protein quantification was performed using BCA protein assay. Total proteins (40 μg/lane) were separated on 4-12% Bis–Tris-polyacrylamide gel electrophoresis (NuPAGE; Thermo Scientific) under reducing conditions and subsequently blotted onto nitrocellulose membranes using iBlot 2 Dry Blotting System (Biorad). Primary antibodies against phospho-Pyk2 Tyr 402 (1:1000; Cell Signalling, cat. no. 3291), total Pyk2 (1:1000; Sigma, cat. no. P3902) and β-actin (1:5000; Abcam, cat. no. ab8226) were used for immunoblotting. After incubation with the appropriate HRP-conjugated secondary antibodies, the protein bands were detected using ImageJ. Samples outside of 3× median absolute deviations were deemed outliers and were excluded from the analysis.

### Statistical analysis

Synapse connectivity data were analyzed using Kruskal-Wallis ANOVA, followed by Wilcoxon rank-sum test to compare individual groups. The statistical unit was microfluidic device. Synapse data were pooled after normalization by the mean of control group for each primary neuron preparation. Human brain data were analyzed using Wilcoxon rank-sum test between cases and controls. Other data were analyzed using unpaired or paired *t*-test as appropriate. A *p*-value less than 0.05 was considered statistically significant.

## Results

### Microfluidic co-culture device to expose hippocampal synapses to synthetic or cell secreted Aβ oligomers

With the goal to better mimic Alzheimer’s disease *in vitro*, we developed a microfluidic device that permits synapse formation between two sets of neurons cultured in distinct chambers. Based on a previous design, our device consists of three distinct chambers, interconnected *via* parallel microchannels that constrain neuronal cell bodies but permit axons and dendrites to cross through (Fig. 1) (Kilinc *et al*., 2014). Adjusting the lengths of microchannels made it possible to allow axons and dendrites from the right (or “postsynaptic”) chamber, but only axons from the left (or “presynaptic”) chamber to reach the central (or “synaptic”) chamber (Fig. 1C) (Taylor *et al*., 2010). As a novel design feature, one end of the synaptic chamber bifurcates, where one branch terminates with an access well and the other one connects to a fourth (co-culture) chamber *via* a dense series of short microchannels. In addition, we employed narrowing microchannels that promote unidirectional neurite crossing (Peyrin *et al*., 2011). When rat postnatal hippocampal neurons were cultured in one chamber only (Supplementary Fig. 3), 2.9× more axons crossed the long microchannels in the forward direction (from the wide end towards the narrow end) than the reverse direction. This figure decreased to 1.9× in the case of short microchannels.

**Figure 1.**
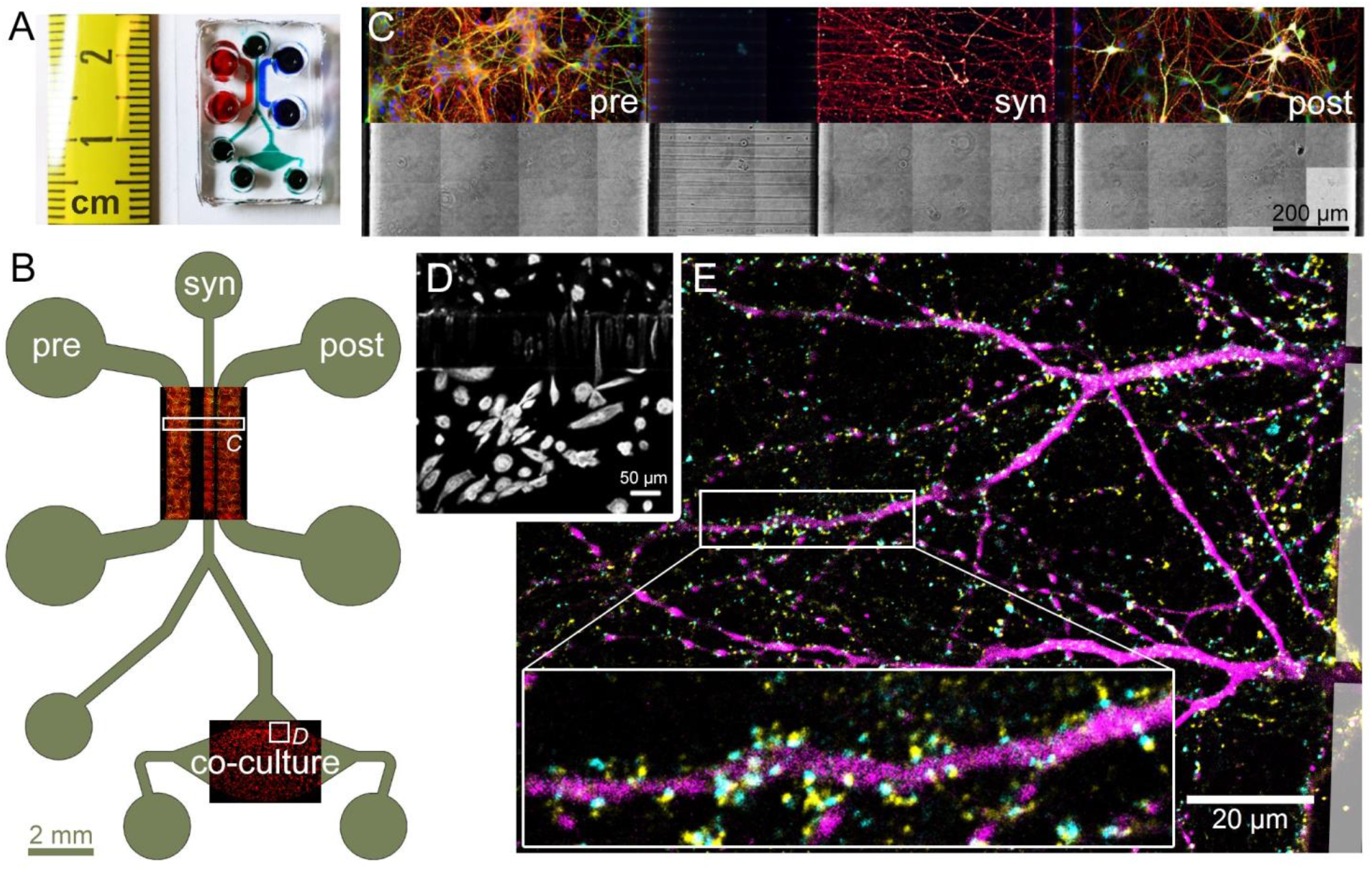
Design and operating principle of the microfluidic co-culture device. **A.** Photograph of the microfluidic device bonded to a coverslip. **B.** The layout of the device showing the presynaptic (pre), synaptic (syn), and postsynaptic (post) chambers, as well as the co-culture chamber housing the CHO cells. Overlays show immunofluorescence images of primary neurons and CHO cells in their respective chambers, stained for α-tubulin (red) and MAP2 (green). **C.** Subcellular compartmentalization of neurons was shown by immunostaining against β3 -tubulin (red) and MAP2 (green), axonal and somatodendritic markers, respectively. Cell bodies were stained with Hoechst (blue). Microchannel structure is evident in the brightfield image of the same area. **D.** 15× magnification of the square marked in (B), showing CHO cells cultured in the co-culture chamber. CHO cells pass through the microchannels, but do not migrate up the synaptic channel. **E.** Synapse formation in the synaptic chamber was evidenced by the localization of Synaptophysin 1 (yellow) and Homer 1 (cyan) puncta, pre- and postsynaptic markers, respectively, around MAP2-positive dendrites (magenta). Boxed area is 3× magnified.

At 14 days *in vitro* (DIV14), no dendrites emanating from neurons cultured in the presynaptic chamber were observed in the synaptic chamber. Axons from these neurons, however, invaded the entire synaptic chamber. 11.0%±4.0% of these axons crossed the short microchannels in the reverse direction and reached the postsynaptic chamber, as measured by the ratio of the β3-tubulin fluorescence between the emitting and receiving chambers. On the other hand, axons and dendrites from neurons cultured in the postsynaptic chamber fully invaded the synaptic chamber by DIV14. None of these dendrites and only 5.9%±1.6% of these axons crossed the long microchannels in the reverse direction and reached the presynaptic chamber. Synapse formation in the synapse chamber was confirmed by immunostaining against Synaptophysin (Syp) 1 and Homer 1, pre- and postsynaptic markers, respectively (Fig. 1E). In summary, the synaptic chamber receives axons from both pre- and postsynaptic chambers, receives dendrites only from the postsynaptic chamber, and contains 83.2%±6.1% of all synaptic connections formed between pre- and postsynaptic chambers.

### Effect of synthetic Aβ_42_ oligomers on synapse connectivity

We oligomerized synthetic Aβ_1-42_ and Aβ_42-1_ (inverted control peptide) and assessed the presence of oligomeric species *via* Coomassie blue staining. We confirmed the presence of low-MW oligomers in the Aβ_1-42_ sample (Fig. 2A). We exposed mature synapses (DIV14) to synthetic Aβ peptides by adding the oligomer solution to the synapse chamber to reach an initial concentration of 100 nM. We kept the media levels in the synaptic reservoirs lower than those in the pre- and postsynaptic reservoirs. The hydrostatic pressure difference induced a flow through the microchannels countering the molecular diffusion, which localized the treatment initially to the synaptic chamber. However, as the pressure-driven flow ceased, the media levels equilibrated and Aβ peptides diffused throughout the device. Since the short microchannels offer little fluidic resistance compared to the long microchannels, it would be safe to assume that the media levels equilibrated first between the synaptic and postsynaptic chambers and then between the synaptic and presynaptic chambers.

**Figure 2.**
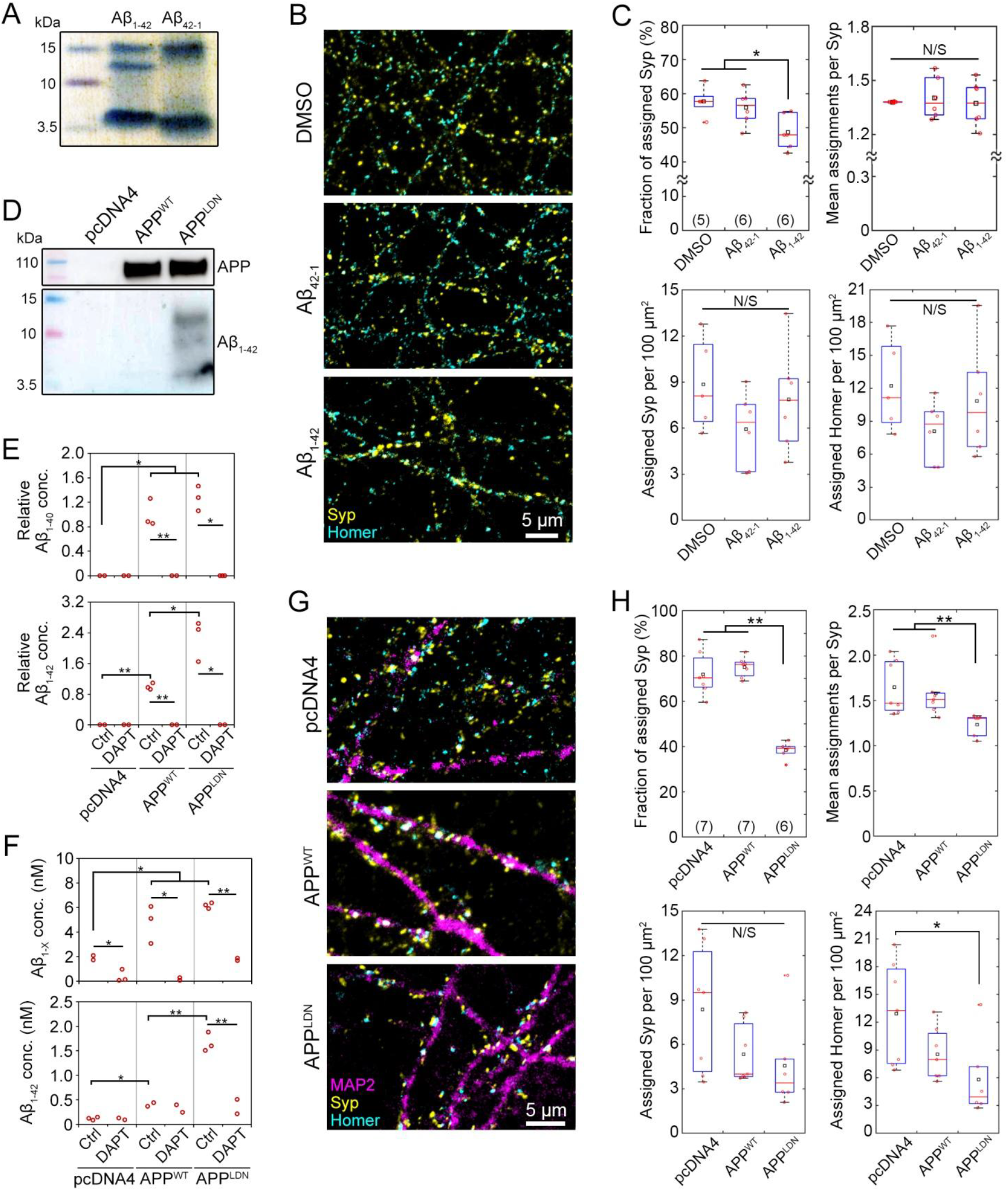
Synthetic and cell-secreted Aβ_1-42_ induce different levels of synaptic toxicity. **A.** Coomassie blue stain of in-house oligomerized synthetic Aβ_1-42_ and Aβ_42-1_ (inverted control) peptides show the presence of low molecular weight oligomers. **B-C.** Exemplary images and synaptic read-outs 16 h after the introduction of synthetic oligomer solution into the synaptic chamber at DIV14 (final concentration = 100 nM). Fraction of Syp puncta assigned by Homer puncta, mean number of Homer puncta assigned per Syp, and densities of assigned Syp and Homer puncta, according to distance-based analysis workflow (see Supplementary Fig. 1 for details). **D.** Immunoblots of CHO cell media two days after stimulation. **E.** Relative Aβ_1-42_ concentration in off-chip CHO cell media quantified *via* ELISA. γ-secretase inhibitor DAPT was applied at 18 μg/mL for 5 d. Data normalized to APP^WT^ control condition. As the sample was concentrated prior to Aβ_1-42_ measurement, Aβ_1-42_ levels cannot be directly compared to Aβ_1-40_ levels. **F.** Alpha-LISA measurement of Aβ_1-X_ and Aβ_1-42_ in media collected from the top well of the synaptic chamber at DIV14. γ-secretase inhibitor DAPT was applied at 18 μg/mL for 5 d. One-way ANOVA, followed by unpaired *t*-test. **G-H.** Exemplary images and synaptic read-outs following co-culture with CHO cells at DIV 14. In box plots, red circles, red bars, black squares, and red plus signs indicate individual data points, sample median and mean, and outliers, respectively. Numbers of microfluidic devices analyzed (obtained from at least 3 independent cultures) are given in parentheses. Analysis based on 416.3 ± 69.2 Syn puncta per image. Kruskal-Wallis ANOVA, followed by Wilcoxon rank-sum test. \**p* < 0.05; ** *p* < 0.005; N/S: not significant.

At the end of the 16 h treatment period, the neurons were fixed and immunostained against pre- and postsynaptic markers. To quantitatively analyze synaptic connectivity, we developed an image analysis workflow based on scanning confocal microscopy, software-assisted identification of pre- and postsynaptic puncta, and proximity-based assignment of postsynaptic puncta to presynaptic puncta (Supplementary Fig. 1). Briefly, each Homer spot was assigned to the nearest Syp spot within a cut-off distance, which was pre-determined using a training set. The fraction of Syp puncta with at least one Homer assignments and the average number of Homer assignments per Syp were determined to be the most robust read-outs of synapse connectivity (see Materials and Methods for details). Exposing synapses to synthetic Aβ_1-42_ oligomers decreased the fraction of assigned Syp without affecting the average number of assignments (Fig. 2B-C). However, the effect size was small (15.6%) and the variation within and among experiments was high.

### Co-culture with CHO cells expressing human APP with London mutation (V717I) induces synaptotoxicity

We cultured Chinese hamster ovary (CHO) cell lines stably overexpressing human APP, either wild-type (CHO-APP^WT^) or with V717I (London) mutation (CHO-APP^LDN^). It has been shown that CHO-APP^WT^ and CHO-APP^LDN^ continuously secrete physiologically-relevant forms of Aβ molecules and CHO-APP^LDN^ provides toxic Aβ species (Guillot-Sestier *et al*., 2012). Immunoblot analysis of media collected from CHO cell cultures confirmed that only the peptides secreted by CHO-APP^LDN^ formed low-MW oligomers (Fig. 2D). In contrast, CHO-pcDNA4 cells, which do not overexpress APP, did not produce any Aβ. We conducted off-chip ELISA measurements to determine the relative levels of Aβ species in CHO cell media (Fig. 2E). As expected, treatment with the γ-secretase inhibitor DAPT completely blocked the secretion of Aβ peptides by both CHO-App^WT^ and CHO-APP^LDN^ cells. Moreover, Äβ_1-42_ (but not Äβ_1-40_) levels in CHO-APP^LDN^ media was higher than in CHO-APP^WT^ media, further supporting the immunoblot results.

To determine the effect of CHO cell-secreted Aβ forms on synapses, we plated *ca*. 10,000 CHO cells in the co-culture chamber 4-6 days prior to the primary neuron culture. This timing was necessary to overcome the problems invoked by the differences of growth media composition between CHO cells and primary neurons. CHO cells proliferated in their growth medium and fully occupied the co-culture chamber. CHO cells were able to cross the short microchannels separating the co-culture chamber from the synaptic chamber (Fig. 1D), but they did not migrate up the synaptic chamber. When the growth medium was replaced with the stimulation medium, the cells stopped proliferating, but continued to secrete Aβ peptides. To confirm that CHO cell-secreted Aβ forms diffused into the synaptic chamber, we collected media from different media reservoirs and quantified their Aβ_1-X_ and Aβ_1-42_ peptide content using corresponding Alpha-LISA kits. Note that media collected from the wells of the microfluidic device did not contain sufficient material for immunoblotting. Measurements taken at DIV14 revealed that the ratio of Aβ_1-42_ to other Aβ forms in the synaptic chamber was 4.4-fold higher in CHO-APP^LDN^ co-cultures than in CHO-APP^WT^ co-cultures (Fig. 2F), as expected for the overexpression of mutated APP (Guillot-Sestier *et al*., 2012). Aβ forms in the media decreased to undetectable levels when CHO cells were treated with γ-secretase inhibitor DAPT for 5 d prior to media collection, further confirming that the majority of the measured Aβ in the synaptic chamber was secreted by the CHO cells. Presence of Aβ_1-X_ in media collected from the synaptic chamber, but not in the co-culture chamber of the CHO-pcDNA4 co-cultures is indicative of neuronal APP processing, since CHO-pcDNA4 cells do not express APP and therefore do not secrete Aβ peptides (Supplementary Fig. 4).

We conducted off-chip experiments to assess the availability of cell-secreted Aβ forms following the addition of the conditioned media to primary neuron cultures. Alpha-LISA measurements conducted at different time points (from 6 h to 7 d) suggested that Aβ concentrations in the media decreased logarithmically over time (Supplementary Fig. 5), suggesting that the peptides degraded or consumed by the cells. In contrast, the concentration of synthetic Aβ_1-42_ peptide did not exhibit such a decrease when added to the primary neuron cultures (Supplementary Fig. 6). These observations further support the idea that cell-secreted Aβ forms need to be periodically resupplied to the neuron culture medium unless a co-culture model is available. When synapses were exposed to CHO cell-secreted Aβ forms for 14 days in the co-culture device, a strong decrease in synapse connectivity was observed in CHO-APP^LDN^, but not in CHO-APP^WT^ co-cultures (Fig. 2G-H). In this case, *i.e*., following chronic exposure, the effect size was large (48.9%) and the variation within and among experiments was low. The average number of Homer puncta assigned per Syp puncta was also significantly lower for CHO-APP^LDN^ co-cultures (22.0% decrease compared to CHO-APP^WT^). To ensure that the observed decrease in synapse connectivity was not a consequence of neuronal cell death due to CHO cell-secreted Aβ forms, we conducted a classical co-culture experiment using cell inserts. No aberrant neuronal cell death was observed at any time point studied, despite the presence of Aβ_1-42_ in the media of CHO-APP^LDN^ co-cultures (Supplementary Fig 7). Separately, MAP2 protein density in images obtained from the synaptic chamber did not vary with the CHO cell type used in co-cultures (Supplementary Fig. 8), showing that the effect of Aβ forms on synapses was not due to an effect on the dendritic network.

### Aβ antibody 3D6 arrests Aβ in the monomeric form and blocks CHO-APP^LDN^-induced synapse loss

To further characterize the CHO cell-secreted Aβ forms, we tested two monoclonal Aβ antibodies, human SAR228810 (8810; Sanofi; 3 μg/mL) and the mouse version of Bapineuzumab (3D6; Janssen; 3 μg/mL). Both antibodies were developed for passive immunotherapy and tested in clinical trials. 8810 antibody targets soluble protofibrillar and fibrillar species of Aβ and is inactive against Aβ monomers and small oligomeric aggregates (Santin *et al*., 2016). In contrast, the 3D6 antibody targets the N-terminal region of Aβ and expected to capture Aβ molecules in the monomeric conformation (Miles *et al*., 2013; Vandenberghe *et al*., 2016).

We conducted off-chip experiments where CHO cells were treated with these antibodies (or with 18 μg/mL DAPT as a negative control) for 5 d. Western blots of the conditioned media showed that DAPT treatment eliminated all Aβ secretion and induced a slight increase in the levels of APP (Fig. 3A). However, WBs of conditioned media following DAPT treatment exhibited a 15 kDa band that overlapped with oligomers of low MW Aβ_1-42_. Treatment with neither antibody affected the presence of low-MW Aβ_1-42_ forms. Interestingly, the 3D6 antibody induced a strong increase in monomeric Aβ_1-42_ levels in both CHO-APP^WT^ and CHO-APP^LDN^ media, in accordance with the idea that 3D6 arrests the peptide in monomeric form and precludes oligomer formation. We also performed Alpha-LISA and ELISA measurements from media collected from the co-culture chamber, following the treatment of CHO cells with the aforementioned antibodies for 5 d. However, the 3D6 antibody interfered with the measurements and data could not be reported.

**Figure 3.**
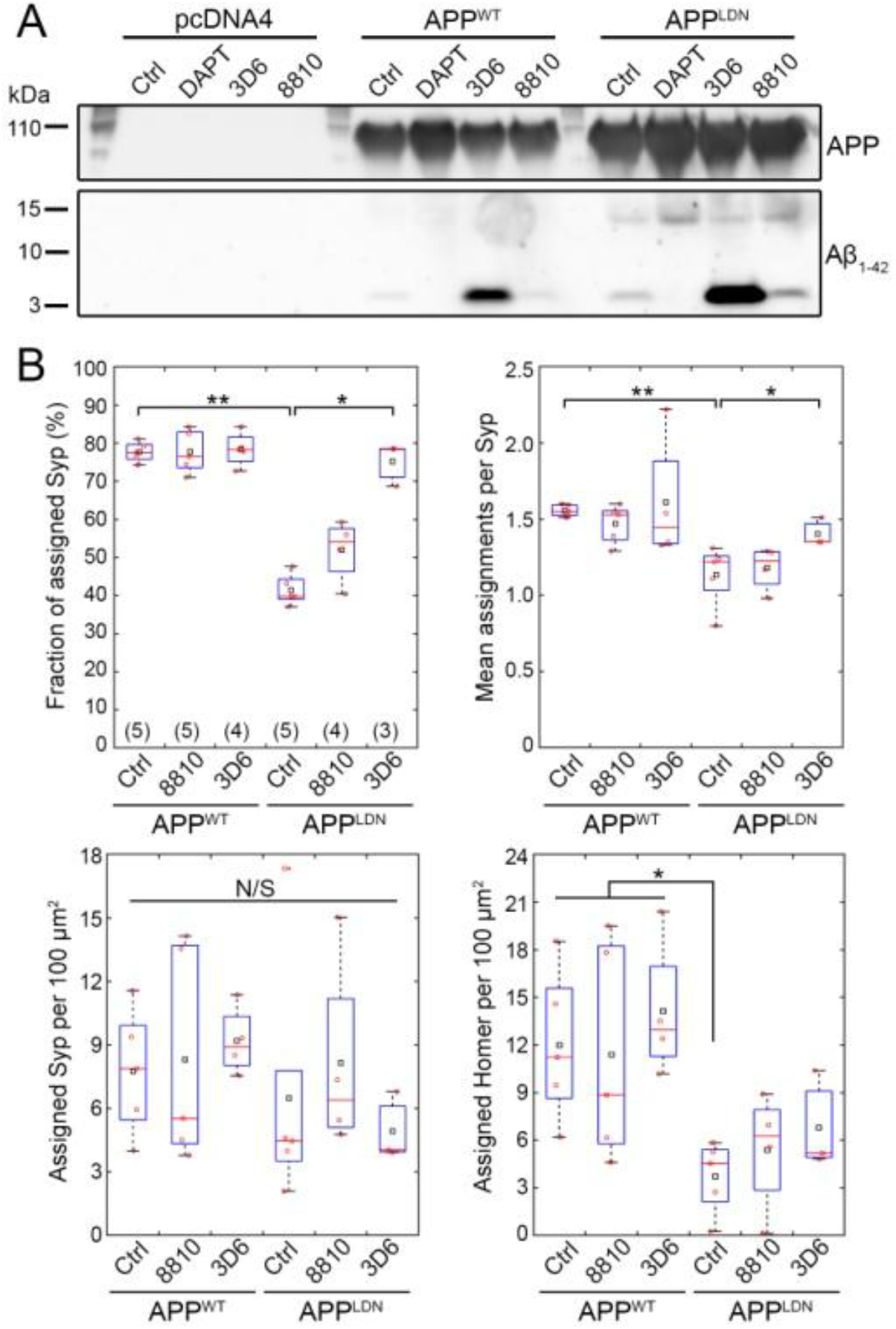
Aβ antibody 3D6 modulates Aβ secretion and blocks the synaptotoxicity due to CHO-APP^LDN^ co-culture. **A.** An exemplary immunoblot of CHO cell media collected after 5 day-long treatment with the indicated compounds (DAPT, 18 μg/mL; antibodies, 3 μg/mL) showing APP cleavage products of different molecular weights. **B.** Synaptic read-outs in antibody-treated co-cultures at DIV14. In box plots, red circles, red bars, and black squares indicate individual data points, sample median, and sample mean, respectively. Numbers of microfluidic devices analyzed (obtained from at least 3 independent cultures) are given in parentheses. Analysis based on 459.5 ± 72.3 Syn puncta per image. Kruskal-Wallis ANOVA, followed by Wilcoxon rank-sum test. * *p* < 0.05; ** *p* < 0.01, N/S: not significant.

We analyzed synaptic connectivity in co-cultures treated with 8810 and 3D6 antibodies for 5 d prior to fixation (Fig. 3B). Similarly to Fig. 2H, untreated CHO-APP^LDN^ co-cultures exhibited a strong (46.6%) decrease in the fraction of Syp puncta assigned by Homer puncta and a significant decrease (26.9%) in the number of Homer assignments per Syp. Treatment with 8810 antibody did not induce a significant difference relative to untreated controls in both cell types. However, treatment with 3D6 antibody completely blocked the effect of CHO-APP^LDN^ on the fraction of Syp assigned and partially blocked the effect of CHO-APP^LDN^ on the number of assignment per Syp. In summary, treatment with an antibody that prevents Aβ monomers from forming oligomers interfered with CHO-APP^LDN^-secreted Aβ species and protected synapses from toxicity likely induced by low-MW oligomers. These findings highlight the potential use of our disease-on-a-chip model and synaptic connectivity analysis for assessing the synapto-protective effects of therapeutic compounds for Alzheimer’s disease, such as Aβ-targeting antibodies.

### Pyk2 overexpression in “postsynaptic” neurons blocks CHO-APP^LDN^-induced decrease in synapse connectivity

We next assessed the relevance of our microfluidic tool to establish whether genetic risk factors of Alzheimer’s disease may be involved in Aβ-dependent synaptotoxicity. Several lines of evidence indicate that genetically driven synaptic failure may occur in Alzheimer’s disease (Dourlen *et al*., 2019) and among the different genetic risk factors susceptible to be studied in our model we focused on *PTK2B* which has been already described to be involved in synaptic functions (Giralt *et al*., 2017).

We evaluated potential changes in the expression of *PTK2B* in the brains of Alzheimer’s disease patients compared to healthy individuals by taking advantage of three publicly available RNAseq datasets: Mayo Clinic data that probed the temporal cortex (Allen *et al*., 2016); ROSMAP data that probed the dorsolateral prefrontal cortex (De Jager *et al*., 2018); and the MSBB data that probed four different brain areas: BA 10, BA 22, BA 36, and BA 44 (Wang *et al*., 2018). This allowed us to investigate potential gene expression changes in brain regions that are affected at different pathological stages of Alzheimer’s disease (Braak and Braak, 1991). We observed a decrease in *PTK2B* expression in Alzheimer’s cases compared to healthy controls in all brain regions analyzed (Table S2); however, after multiple testing correction this decrease was significant only in the BA 22 (16.69% decrease; *p*_adj_ = 1.99×10^-02^), BA 36 (28.89% decrease; *p*_adj_ = 8.49×10^-05^), and BA 44 (15.18% decrease; *p*_adj_ = 3.87×10^-02^) regions of the MSBB dataset, as well as in the ROSMAP dataset (28.89% decrease; *p*_ad_j = 4.57×10^-03^). Consistent with the RNAseq data, we observed a decreasing trend in Pyk2 total protein levels and an increasing trend in the p-Pyk2 protein levels in the brains of Alzheimer’s patients compared to healthy controls (Supplementary Fig. 9). These changes resulted in a significant increase in the relative phosphorylation of Pyk2 in the hippocampus (4.1-fold increase in the p-Pyk2:Pyk2 ratio; p = 0.0274; Wilcoxon rank sum test; 17 cases vs. six controls) and in the cortex (3.6-fold increase in the p-Pyk2:Pyk2 ratio; p = 0.0274; Wilcoxon rank sum test; 17 cases vs. seven controls), suggesting a compensatory mechanism.

We then evaluated the levels of Pyk2 and phospho-Pyk2 in primary neurons as a function of Aβ exposure. We conducted off-chip synaptosome extraction using primary cortical neurons treated with CHO cell media and observed total Pyk2 and p-Pyk2 in both non-PSD and PSD fractions. Relative to CHO-APP^WT^ medium, CHO-App^LDN^ medium caused significant decreases in the relative amounts of total Pyk2 and p-Pyk2 in the postsynaptic fraction (Fig. 4A). In separate experiments based on hippocampal neurons cultured in microfluidic devices, we immunostained p-Pyk2 (Tyr402) alongside synaptic markers and observed that signals localized to both pre- and postsynaptic puncta, with an increased tendency towards the latter (Fig. 4B). We extended our distance-based synaptic connectivity analysis to quantitatively analyze the distribution of p-Pyk2 signals relative to identified synapses, *i.e*., Syp–Homer pairs. Our data showed that 1.5-fold more p-Pyk2 puncta were localized near postsynaptic puncta than near presynaptic puncta regardless of co-culture with CHO-APP^WT^ or CHO-APP^LDN^ cells (Supplementary Fig. 10).

**Figure 4.**
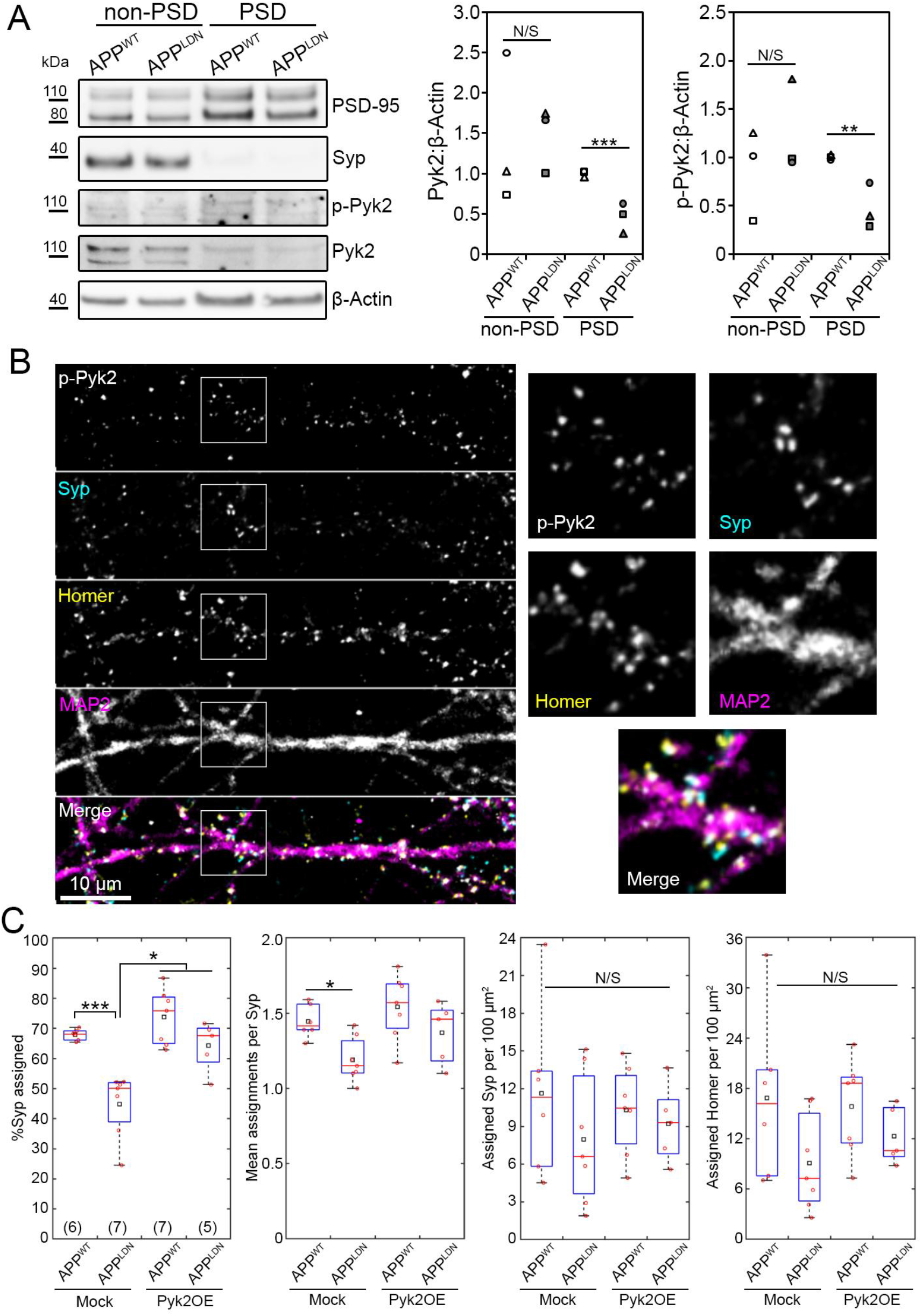
Pyk2 overexpression in postsynaptic neurons blocks the synaptotoxicity due to CHO-APP^LDN^ co-culture. **A.** Exemplary immunoblots and quantification of the postsynaptic (PSD) fraction following synaptosome extraction at DIV14 following 16 h-long treatment with the indicated CHO cell medium. N = 3 independent experiments. Unpaired *t*-test. **B.** An exemplary immunofluorescence image of the synaptic chamber at DIV14 showing phospho-Pyk2 Tyr402 (p-Pyk2), Synaptophysin 1 (Syp) Homer 1, and MAP2. Boxed areas are 2.5× magnified. **C.** Synaptic read-outs following Pyk2 overexpression in postsynaptic neurons from DIV7 to DIV14. In box plots, red circles, red bars, and black squares indicate individual data points, sample median, and sample mean, respectively. Numbers of microfluidic devices analyzed (obtained from at least 3 independent cultures) are given in parentheses. Kruskal-Wallis ANOVA, followed by Wilcoxon rank-sum test. Error bars = SEM. * *p* < 0.05; ** *p* < 0.01; *** p < 0.005; N/S: not significant.

Since we observed (i) a decrease in *PTK2B* expression and a decreasing trend in Pyk2 protein levels in Alzheimer’s brains, (ii) a decrease in Pyk2 levels in the PSD fraction of cortical neurons upon treatment with CHO-APP^LDN^ media, and (iii) a decrease in synaptic connectivity in hippocampal neurons upon co-culture with CHO-APP^LDN^ cells, we hypothesized that Pyk2 was protective and its overexpression could rescue the detrimental effect of CHO-APP^LDN^ co-culture on synapses. Since the active form of Pyk2 was strongly associated with postsynapses and CHO-APP^LDN^ medium affected Pyk2 levels specifically in the PSD fraction, we decided to take advantage of microfluidic compartmentalization and modulate Pyk2 expression in postsynaptic neurons. To this end, we first verified the overexpression of Pyk2 off-chip *via* lentiviral transduction of the relevant cDNA (Supplementary Fig. 11). Next, by using lentiviruses expressing fluorescent proteins, we confirmed that the viral transduction in the microfluidic device was restricted to the target chamber (selectively to pre- or postsynaptic chambers; Supplementary Fig. 12). Over-expressing Pyk2 in the postsynaptic chamber blocked the detrimental effect of CHO-APP^LDN^ co-culture on synaptic connectivity, as evidenced by 12.9% decrease in the fraction of Syp puncta assigned by Homer puncta (as compared to 33.9% decrease when overexpressing the control vector; Fig. 4C). As expected, synaptic connectivity in CHO-APP^WT^ co-cultures was not affected by Pyk2 overexpression (8.7% increase in the fraction of Syp assigned by Homer), confirming that the synaptoprotective effect of postsynaptic Pyk2 overexpression was specific to Aβ toxicity due to CHO-APP^LDN^ co-culture.

## Discussion

The use of microfluidic culture devices for isolating synapses from neuronal cell bodies has been previously demonstrated (Taylor *et al*., 2010; Virlogeux *et al*., 2018). Our co-culture device combines two design concepts: First, by employing different microchannel lengths, it guarantees that dendrites from one but not the other neuron chamber can arrive to the synapse chamber. Second, it minimizes the penetration of axons from the synapse chamber to the neuron chambers and thereby facilitates the passage of axons in the intended direction: 2.9 and 1.9-fold more axons crossed the microchannels in the intended direction relative to the reverse direction, respectively for long and short microchannels. These ratios are indicative of a limited effect of the narrowing channel design on hippocampal neurons and are consistent with earlier reports (Peyrin *et al*., 2011). Our co-culture device also facilitated the concentration of synapses in the synapse chamber: More than 83% of all synapses formed between neurons cultured in the pre- and postsynaptic chambers were found in the synapse chamber. It is important to note that while all dendrites in the synapse chamber arrive from the postsynaptic chamber, the opposite statement is not true, *i.e*., not all axons in the synapse chamber arrive from the presynaptic chamber. Thus, for all synapses analyzed, overexpressing Pyk2 in the postsynaptic chamber guarantees Pyk2 overexpression in the postsynaptic neuron (provided that it is infected by the lentivirus); however, the possibility of Pyk2 overexpression also in the presynaptic neuron cannot be ruled out. However, this uncertainty is not relevant for studies focusing solely on postsynaptic mechanisms. The microfluidic approach provides a simple yet robust method to image and analyze synapses independently of densely-plated cell bodies. Although not exploited in the context of this study, our co-culture device would also permit acute treatment of synapses independently of their cell bodies and independently of the co-cultured cells, thanks to the additional access well connected to the synaptic chamber (Fig. 1B). However, long-term treatments, as in the case for Aβ molecules secreted by the CHO cells, cannot be exclusively directed to synapses due to molecular diffusion.

Several parameters can be considered when inducing Aβ-dependent synaptotoxicity: (i) use of synthetic versus organic oligomers; (ii) acute versus chronic treatment. Our microfludic co-culture model combines organic oligomers at physiological concentrations with chronic treatments and consistently induces Aβ-dependent synaptotoxicity. This is in agreement with a recent report that synthetic oligomers do not assume the same molecular structure as organic oligomers (Kollmer *et al*., 2019), suggesting potential differences in their biological effects. Interestingly, as opposed to synthetic Aβ oligomers, organic Aβ appears to be degraded in our primary neuronal cultures, further highlighting the difference in bioavailability between the two. This observation also indicates that the co-culture model is better adapted for synaptotoxicity studies since it maintains physiological concentrations thanks to the continuous secretion of Aβ peptides from CHO cells. This allows for the analysis of synapse connectivity in response to chronic exposure of synapses to organic Aβ oligomers, as opposed to acute exposure to conditioned media, for instance.

It is important to note that the Aβ peptides secreted by the CHO-APP^LDN^ cells do diffuse into the various chambers of the microfluidic device, considering the long duration of co-culture experiments. Thus, the synaptic toxicity observed could be due to a local effect on synapses or due to an effect on neuronal cell bodies. Through conventional co-culture experiments conducted off-chip we showed that CHO cell-secreted Aβ forms do not induce neuronal death; however, this does not allow us to rule out any potential mechanisms originating from neuronal cell bodies and resulting in the decrease of synaptic connectivity observed. If true, such a mechanism would be more likely to occur in neurons in the postsynaptic chamber, as the fluidic barrier between the synaptic and postsynaptic chambers is much weaker than that between the synaptic and presynaptic chambers, allowing the cell secreted Aβ to easily access the postsynaptic chamber.

The role of Pyk2 in synapses appears to be complex, considering the seemingly opposite results of recent studies: Pyk2 has been shown to be required for long-term potentiation (LTP) (Huang *et al*., 2001). However, others have shown that Pyk2 is not required for LTP, but for long-term depression (Hsin *et al*., 2010; Salazar *et al*., 2019), and that Pyk2 overexpression inhibits LTP (Hsin *et al*., 2010) and induces dendritic spine loss (Lee *et al*., 2019). Recent analysis of protein synthesis and degradation during synaptic scaling showed that Pyk2 protein level was significantly increased in response to drug-induced decrease in network activity (*via* increased synthesis and decreased degradation) (Dörrbaum *et al*., 2020). The role of Pyk2 in Aβ toxicity has also been debated, where it has been shown to be deleterious (Salazar *et al*., 2019) or protective (Giralt *et al*., 2018) *in vivo*. Our findings are in agreement with the notion that Pyk2 localizes to the postsynaptic compartment (Giralt *et al*., 2017; Lee *et al*., 2019; Salazar *et al*., 2019). To shed further light into the role of Pyk2 in Aβ-induced synapse toxicity we overexpressed Pyk2 specifically in postsynaptic neurons, since we observed that the active form of Pyk2 was strongly associated with postsynapses and that Aβ treatment affected Pyk2 levels specifically in the PSD fraction. At the pathophysiological level, several of our observations fit well with the results of recent *in vivo* studies using Alzheimer’s disease-like mouse models: (i) decreased Pyk2 activity has been reported in 5xFAD mice (Giralt *et al*., 2018); (ii) rescue of Pyk2 expression improved the behavioral and synaptic molecular phenotypes of the double transgenic 5xFAD×Pyk^-/-^ mouse model (Giralt *et al*., 2018). Interestingly, this rescue seems to have no impact on Aβ loads, suggesting that Pyk2 overexpression impacts pathophysiological processes downstream of Aβ production and amyloid deposition. This observation is in agreement with our results suggesting that Pyk2 overexpression in postsynaptic neurons may restrict Aβ-induced synaptotoxicity. In stark contrast to these reports, increased Pyk2 activation has been shown in response to acute Aβ oligomer treatment in brain slices (Haas *et al*., 2016; Haas and Strittmatter, 2016) and in the APP/PS1 mouse model (Kaufman *et al*., 2015). Moreover, Pyk2 has been shown to be detrimental in the double transgenic APP/PS1×Pyk^-/-^ mouse model, where lack of Pyk2 protected from synapse loss and memory impairment (Salazar *et al*., 2019). These contradictory reports may be due to differences in the disease model and in the Pyk2^-/-^ model used (Giralt *et al*., 2017; Salazar *et al*., 2019). In summary, the sparse literature on Pyk2’s role in synapses is highly controversial and calls for further *in vitro* and *in vivo* work.

In conclusion, our microfluidic co-culture device provides an *in vitro* model of Aβ synaptotoxicity based on exposing synapses of primary hippocampal neurons to cell-secreted Aβ_1-42_ peptides. This disease-on-a-chip model is highly relevant to Alzheimer’s disease in several aspects: (i) long-term, low-dose exposure to organic Aβ forms is preferable over acute treatments with synthetic oligomers at high concentrations; (ii) isolating synapses in a separate microfluidic chamber facilitates the analysis of synaptic connectivity *via* immunostaining pre- and postsynaptic markers without the interference of cell bodies; (iii) providing exclusive access to neurons cultured in the pre- and postsynaptic chambers to selectively under- or overexpress Alzheimer’s disease genetic risk factors therein may potentially help dissect their pre- and postsynaptic roles. Deciphering the mechanisms by which the genetic risk factors contribute to Alzheimer’s pathology may lead to novel therapeutic approaches.

## Acknowledgements

This work was partly supported by the French RENATECH network (P-16-01891). This study was funded by INSERM, Institut Pasteur de Lille, the EU Joint Programme – Neurodegenerative Diseases Research (JPND; 3DMiniBrain), Agence Nationale de la Recherche (ANR-19-CE16-0020), and Fondation Vaincre Alzheimer (FR-17006p). This work was also funded by the Lille Métropole Communauté Urbaine and the French government’s LABEX DISTALZ program (Development of innovative strategies for a transdisciplinary approach to Alzheimer’s disease). D. M.-C. was supported by a PhD scholarship from Coordenação de Aperfeiçoamento de Pessoal de Nível Superior (CAPES). This work was also co-funded by the European Union under the European Regional Development Fund (ERDF) and by the Hauts de France Regional Council (contract no. 18006176), the Métropole Européenne de Lille (contract no. 2016_ESR_05), and the French State (contract no. 2018-3-CTRL_IPL_Phase2). The aforementioned funding bodies did not play any roles in the design of the study and collection, analysis, and interpretation of data and in writing the manuscript.

The authors thank the BICeL platform of the Institut Biologie de Lille. The authors thank Laurent Pradier and Philippe Bertrand at Sanofi for fruitful discussions. The authors thank Karine Blary at the IEMN Lille for the microfabrication work. The authors thank the vectorology platform Transbiomed for lentivirus production. The authors thank Charles Duyckaerts and the “NeuroCEB” Brain Bank (GIE Neuro-CEB BB-0033-00011) for providing the brain tissue samples.

The results published here are in whole or in part based on data obtained from the AMP-AD Knowledge Portal (https://adknowledgeportal.synapse.org/).

These data were generated from postmortem brain tissue collected through the Mount Sinai VA Medical Center Brain Bank and were provided by Dr. Eric Schadt from Mount Sinai School of Medicine.

Study data were also provided by the following sources: The Mayo Clinic Alzheimers Disease Genetic Studies, led by Dr. Nilufer Taner and Dr. Steven G. Younkin, Mayo Clinic, Jacksonville, FL using samples from the Mayo Clinic Study of Aging, the Mayo Clinic Alzheimers Disease Research Center, and the Mayo Clinic Brain Bank. Data collection was supported through funding by NIA grants P50 AG016574, R01 AG032990, U01 AG046139, R01 AG018023, U01 AG006576, U01 AG006786, R01 AG025711, R01 AG017216, R01 AG003949, NINDS grant R01 NS080820, CurePSP Foundation, and support from Mayo Foundation. Study data includes samples collected through the Sun Health Research Institute Brain and Body Donation Program of Sun City, Arizona. The Brain and Body Donation Program is supported by the National Institute of Neurological Disorders and Stroke (U24 NS072026 National Brain and Tissue Resource for Parkinsons Disease and Related Disorders), the National Institute on Aging (P30 AG19610 Arizona Alzheimers Disease Core Center), the Arizona Department of Health Services (contract 211002, Arizona Alzheimers Research Center), the Arizona Biomedical Research Commission (contracts 4001, 0011, 05-901 and 1001 to the Arizona Parkinson’s Disease Consortium) and the Michael J. Fox Foundation for Parkinsons Research.

Study data were also provided by the Rush Alzheimer’s Disease Center, Rush University Medical Center, Chicago. Data collection was supported through funding by NIA grants P30AG10161 (ROS), R01AG15819 (ROSMAP; genomics and RNAseq), R01AG17917 (MAP), R01AG30146, R01AG36042 (5hC methylation, ATACseq), RC2AG036547 (H3K9Ac), R01AG36836 (RNAseq), R01AG48015 (monocyte RNAseq) RF1AG57473 (single nucleus RNAseq), U01AG32984 (genomic and whole exome sequencing), U01AG46152 (ROSMAP AMP-AD, targeted proteomics), U01AG46161(TMT proteomics), U01AG61356 (whole genome sequencing, targeted proteomics, ROSMAP AMP-AD), the Illinois Department of Public Health (ROSMAP), and the Translational Genomics Research Institute (genomic). Additional phenotypic data can be requested at www.radc.rush.edu.

---

**Table S1.** Demographic details of the neuropathological cohort. ND: Not determined. AD: Alzheimer’s disease.

| *Individual* | *Braak stage* | *Gender* | *Age at death* | *Post-mortem delay (h)* | *Neuropathological diagnosis* |
| --- | --- | --- | --- | --- | --- |
| 1 | III | F | 95 | ND | Non-AD |
| 2 | – | F | 52 | 29 | Non-AD |
| 3 | I | F | 92 | 21 | Non-AD |
| 4 | II | M | 82 | 63 | Non-AD |
| 5 | II | F | 83 | 21 | Non-AD |
| 6 | III | F | 93 | 24 | Non-AD |
| 7 | – | M | 69 | 6 | Non-AD |
| 8 | IV | F | 76 | 28 | Non-AD |
| 9 | VI | F | 48 | ND | AD |
| 10 | VI | M | 57 | 19 | AD |
| 11 | VI | F | 60 | ND | AD |
| 12 | VI | M | 72 | 44 | AD |
| 13 | VI | F | 72 | 5 | AD |
| 14 | VI | F | 78 | 24 | AD |
| 15 | VI | M | 81 | 17 | AD |
| 16 | VI | F | 85 | 31 | AD |
| 17 | VI | F | 100 | 67 | AD |
| 18 | VI | M | 67 | 30 | AD |
| 19 | VI | M | 67 | ND | AD |
| 20 | VI | M | 53 | ND | AD |
| 21 | VI | F | 56 | 26 | AD |
| 22 | VI | F | 70 | ND | AD |
| 23 | VI | F | 72 | 27 | AD |
| 24 | VI | M | 76 | 27 | AD |
| 25 | VI | F | 77 | 48 | AD |
| 26 | VI | F | 81 | 41 | AD |
| 27 | VI | F | 85 | 28 | AD |
| 28 | VI | F | 90 | 24 | AD |

**Table S2.**
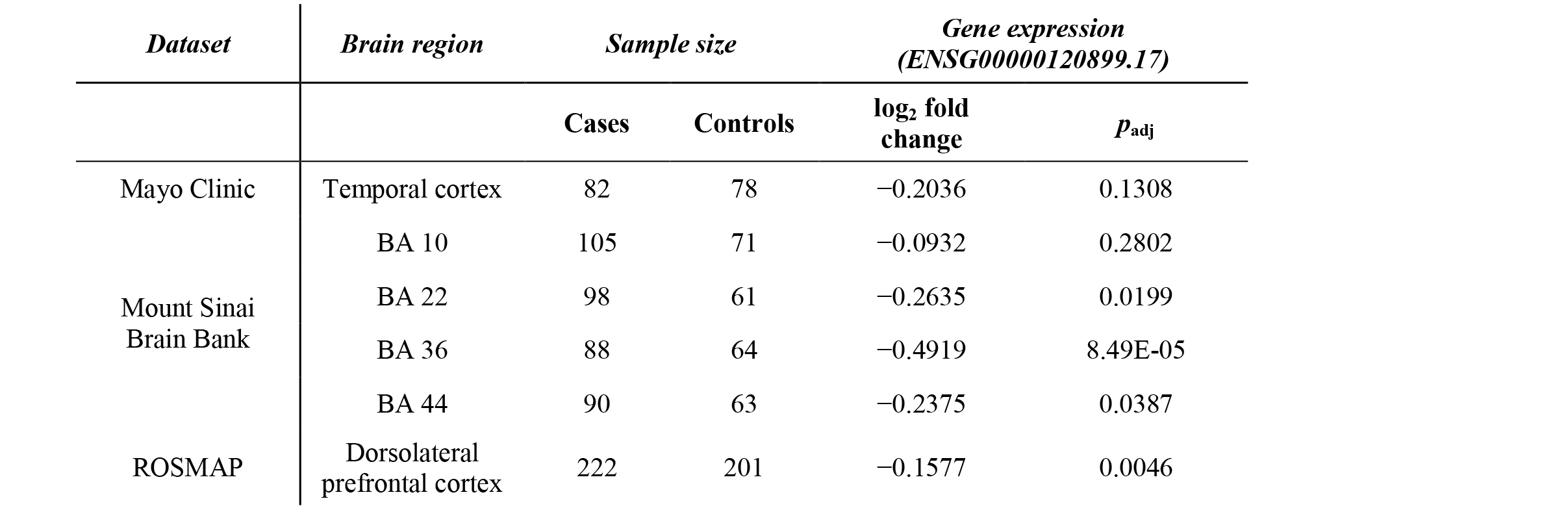
*PTK2B* gene expression analysis in different publicly-available datasets

**Figure S1.**
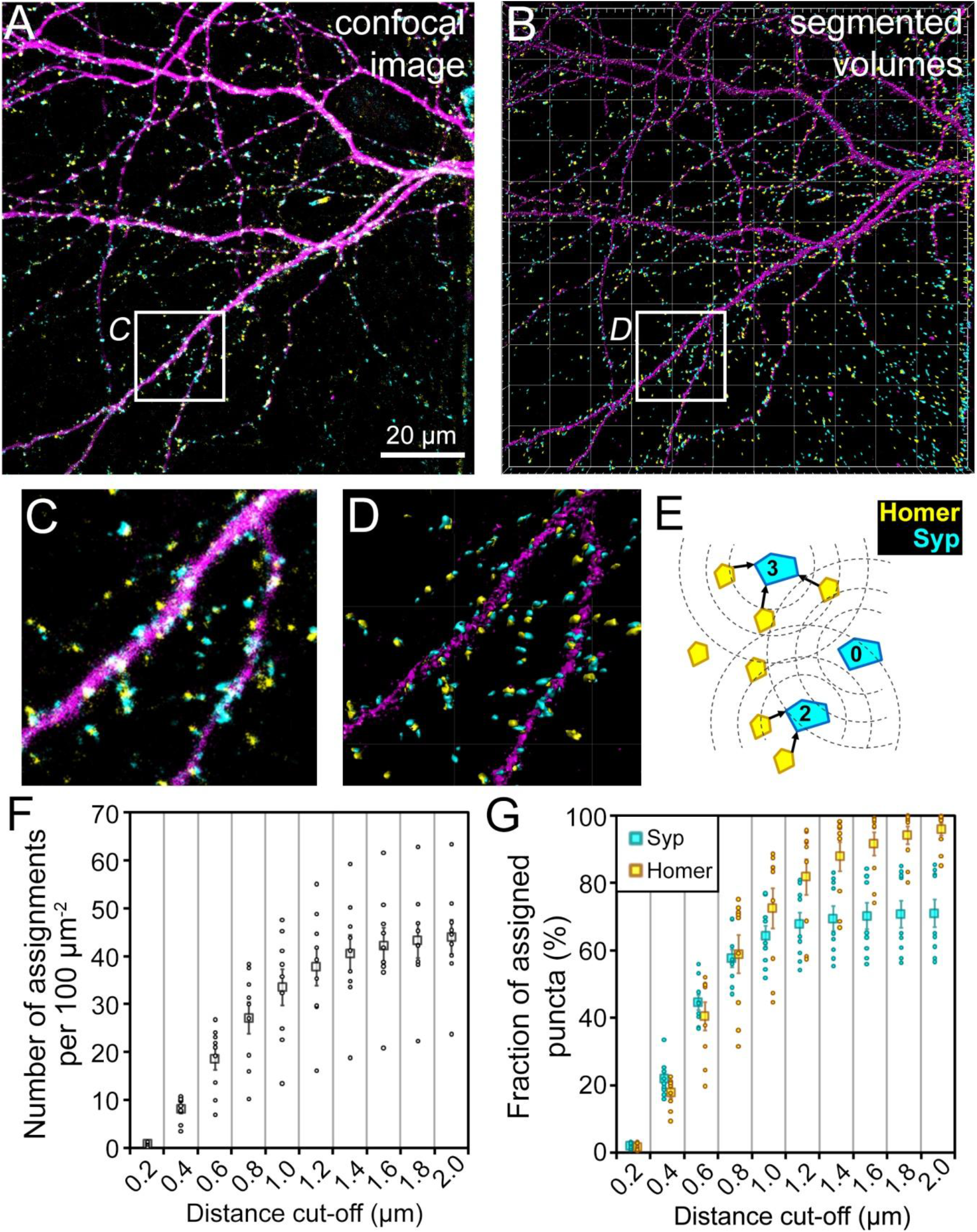
Synapse analysis workflow and the determination of the distance cut-off. **A.** An exemplary dendrite in the synaptic chamber. Image shows the maximum intensity projection of Homer 1 (yellow), Synaptophysin 1 (cyan), and MAP2 (magenta) stains. **B.** Segmentation of the same image in Imaris. **C-D.** Marked areas in *A* and *B*, respectively, magnified 3.5×. **E.** Principle of assigning postsynaptic (Homer) to presynaptic (Syp) puncta based on proximity. Numbers indicate the number of assignments for each presynaptic puncta. **F.** Synaptic assignment density as a function of distance cut-off. **G.** Fractions of assigned pre- and postsynaptic puncta as a function of distance cut-off, showing that Syp assignments saturate at around 1.0 μm for the dataset analyzed. Data in *F* and *G* are based on N = 9 microfluidic devices (small circles) from 4 independent cultures; 5-8 images per device. Rectangles and error bars show show sample mean and SEM, respectively.

**Figure S2.**
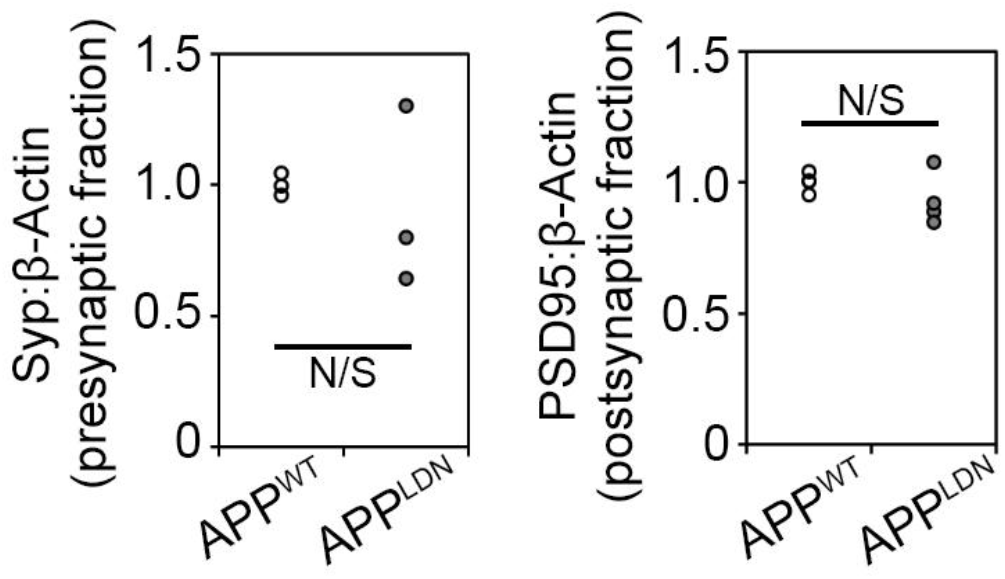
Quantification of synaptic markers in the synaptosomal fraction. Synaptophysin (Syp) in the presynaptic fraction and PSD95 in the postsynaptic fraction for primary neuronal cultures exposed to media collected from CHO-APP^WT^ and CHO-APP^LDN^ cultures. Each data point represents one independent experiment. Unpaired *t*-test. N/S: not significant.

**Figure S3.**
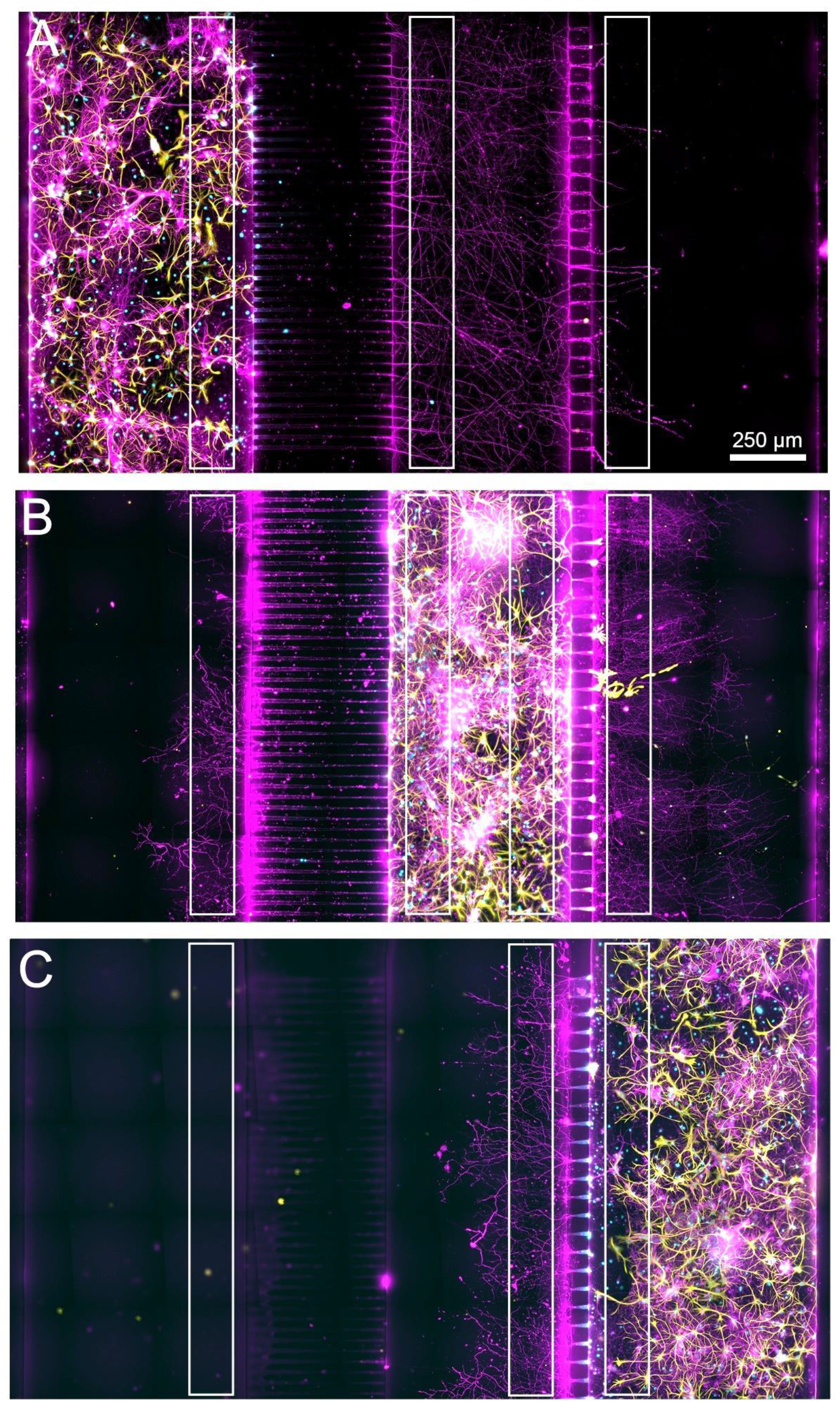
Exemplary images of microfluidic devices where primary neurons were plated only in the presynaptic (**A**), synaptic (**B**), or the postsynaptic (**C**) chamber. β3-tubulin (magenta), Hoechst (cyan), and GFAP (yellow) staining shown at DIV14. White rectangles indicate regions where β3-tubulin staining intensity was measured to calculate the penetration ratios.

**Figure S4.**
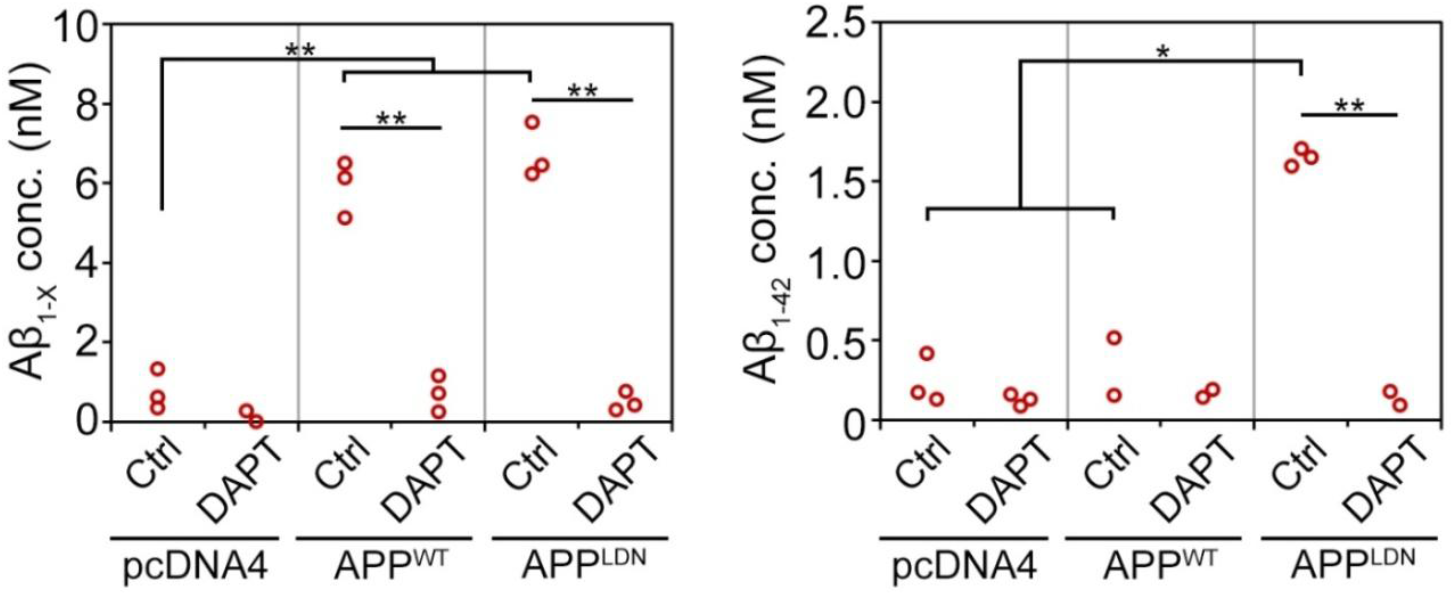
Concentration of Aβ_1-X_ and Aβ_1-42_ in the media collected from the reservoir of the co-culture chamber at DIV14 measured *via* Alpha-LISA. γ-secretase inhibitor DAPT was applied at 18 μg/mL for 5 d. One-way ANOVA, followed by unpaired *t*-test. * *p* < 0.05; ** *p* < 0.005.

**Figure S5.**
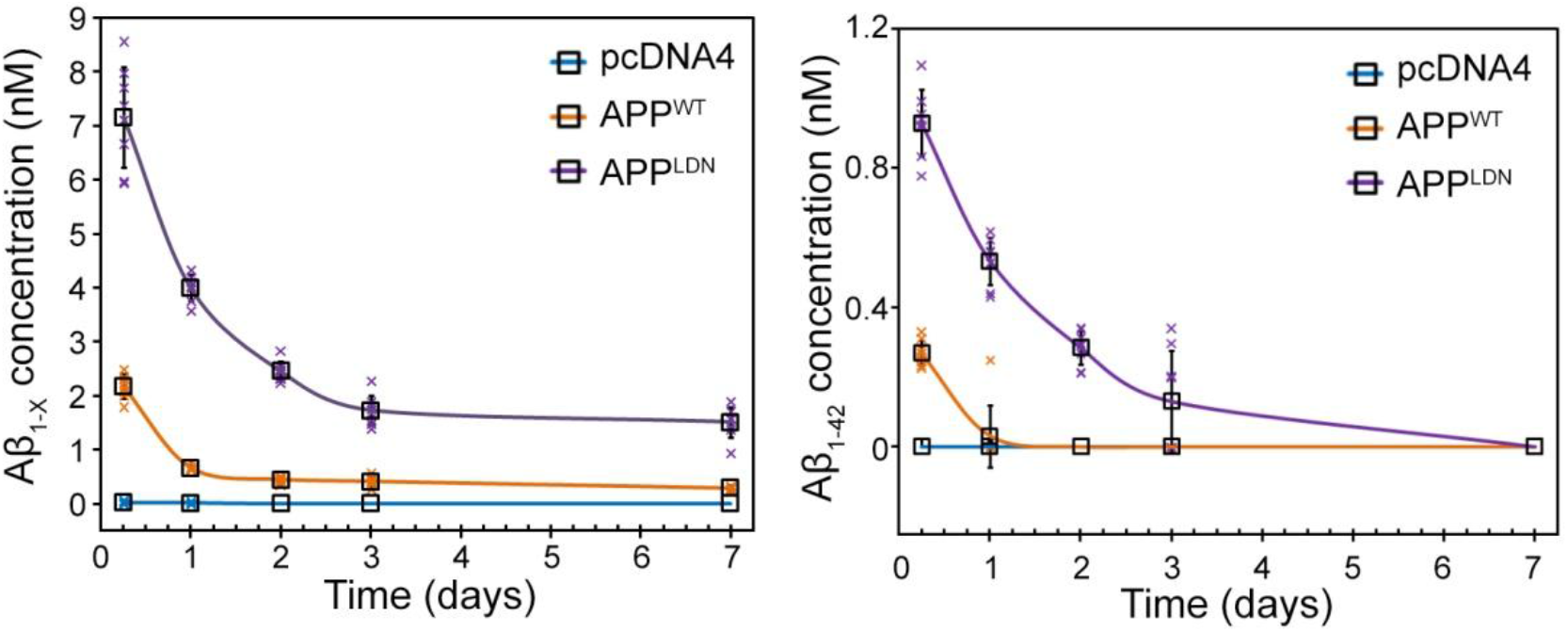
Concentration of Aβ_1-X_ and Aβ_1-42_ in the media collected from primary neurons cultured in 384-well plates and exposed to CHO cell-conditioned media for 6 h to 7 d. Concentration was determined *via* Alpha-LISA using 6-8 wells per condition. Error bars indicate standard deviation of the mean. Individual data points are not shown for time points where all values were equal to zero.

**Figure S6.**
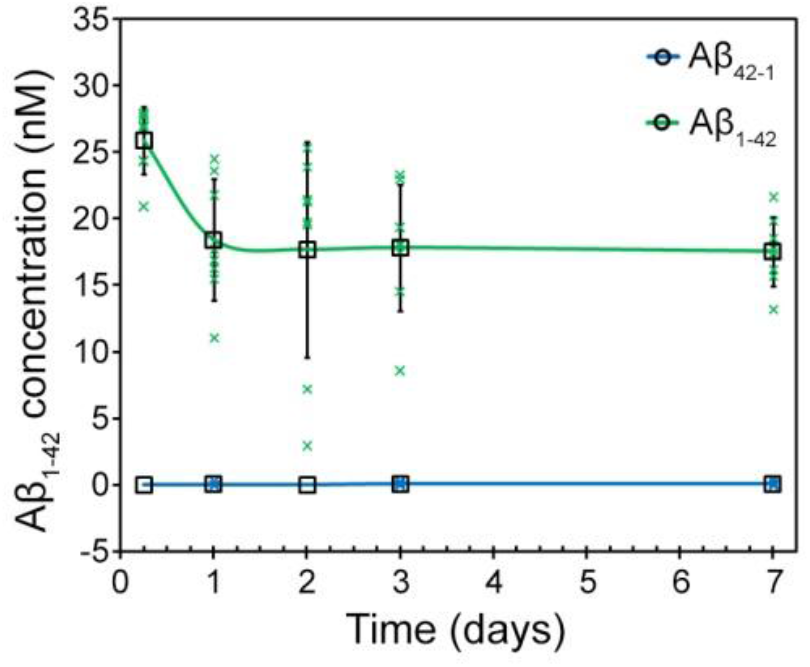
Concentration of Aβ_1-42_ in the media collected from primary neurons cultured in 384-well plates and treated with synthetic Aβ_1-42_ oligomers for 6 h to 7 d. Concentration was determined *via* Alpha-LISA using 6-8 wells per condition. Error bars indicate standard deviation of the mean. Individual data points are not shown for time points where all values were equal to zero.

**Figure S7.**
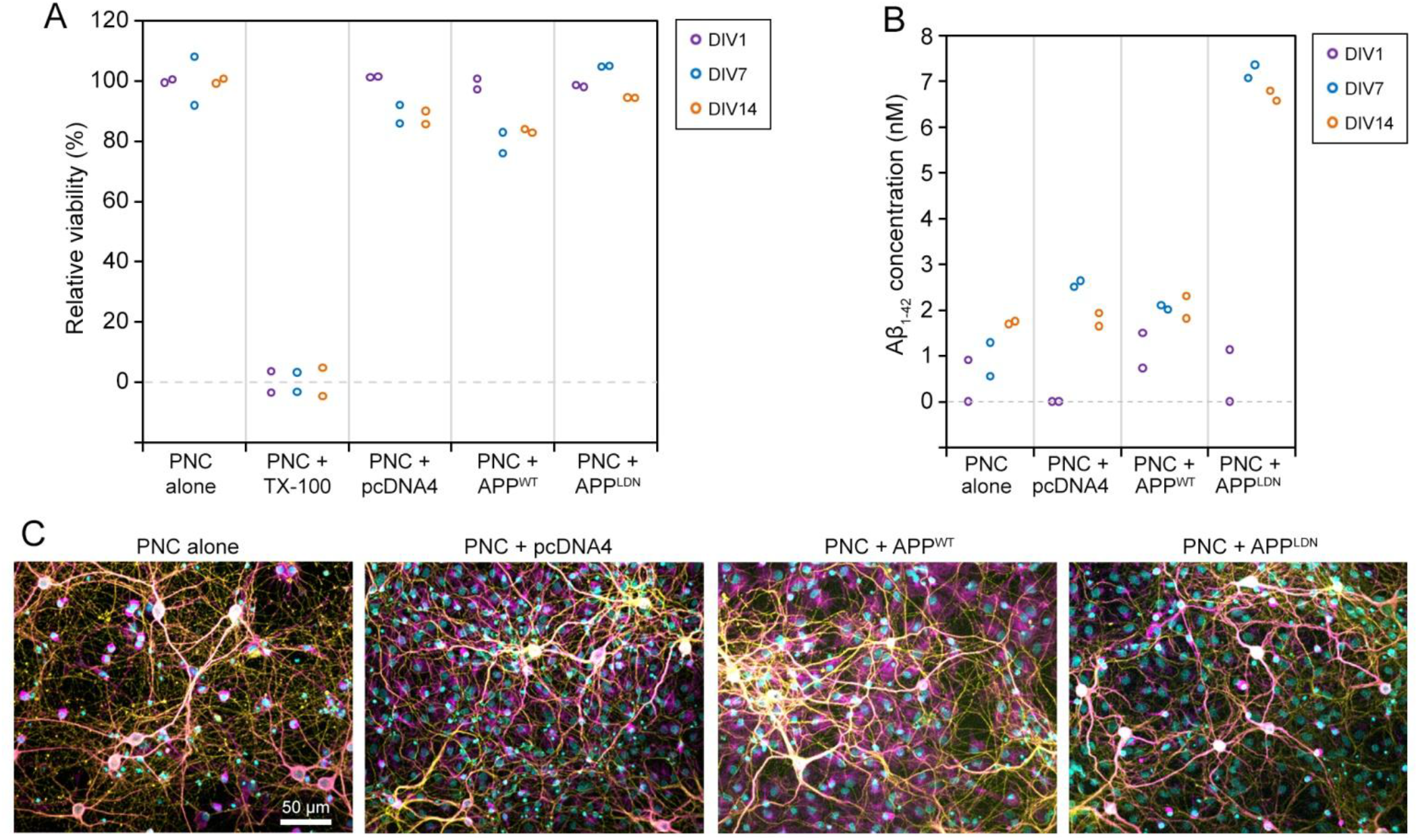
Cell viability in conventional co-cultures. **A.** LDH release assay in 24-well plates with Transwell inserts. Primary neuronal culture (PNC) alone and PNC treated with 1% Triton X-100 (TX-100) were used as positive (100%) and negative (0%) controls, respectively, to calculate the relative cell viability, separately for each time point. **B.** Aβ_1-42_ concentration in the mono- and co-culture media measured *via* Alpha-LISA. Each dot refers to a single well of the 24-well plate. Measurements below detection limit were shown as zero in panel B. **C.** Exemplary images of neurons from the same assay, showing MAP2 (magenta), β3-tubulin (yellow), and Hoechst (cyan) staining.

**Figure S8.**
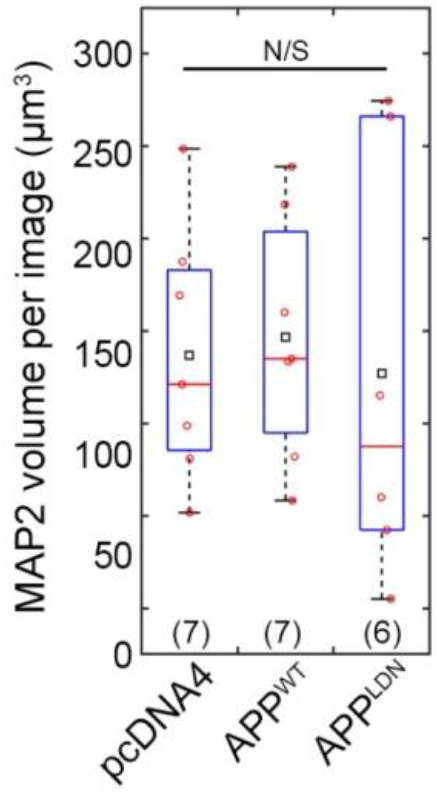
MAP2 levels in the synaptic chamber following co-culture with CHO cells at DIV 14. MAP2 volumes were detected in Imaris. See Fig. 2G for synaptic read-outs obtained from the same set of images. In the box plot, red circles, red bars and black squares indicate individual data points, sample median, and sample mean, respectively. Numbers of microfluidic devices analyzed (obtained from at least 3 independent cultures) are given in parentheses. Kruskal-Wallis ANOVA. N/S: not significant.

**Figure S9.**
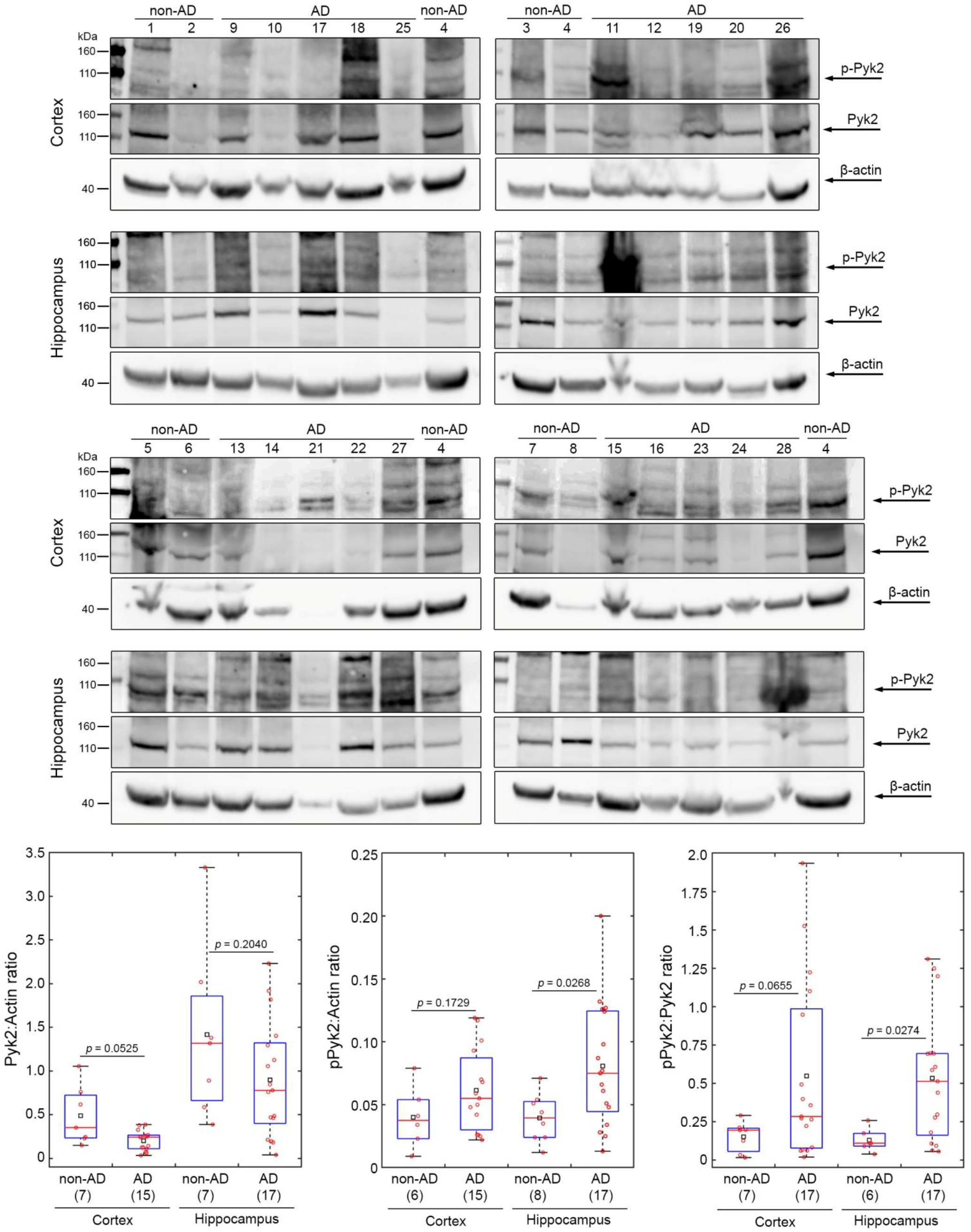
Total and phospho-Pyk2 levels in the post-mortem brains of Alzheimer’s disease cases and healthy controls. Demographic details of the cohort are given in Table S1. In box plots, red circles, red bars, and black squares indicate individual data points, sample median and mean, respectively. Numbers of individual data points analyzed for each condition are given in parentheses. Wilcoxon rank-sum test.

**Figure S10.**
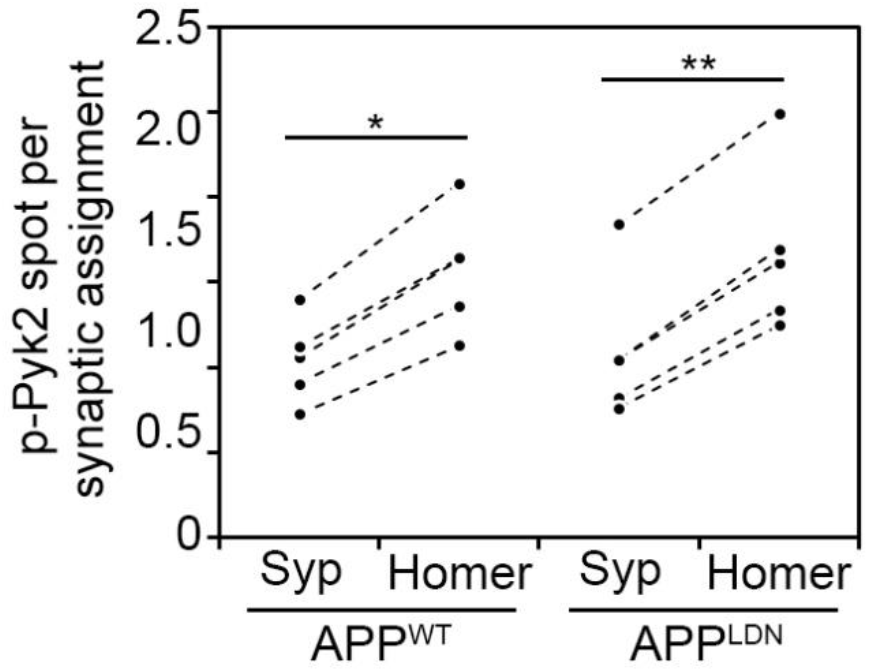
Localization of phospho-Pyk2 Tyr402 (p-Pyk2) puncta relative to synapses. Each Homer spot was assigned to the nearest Synaptophysin (Syp) spot within cut-off distance of 1.0 μm. Each p-Pyk2 spot was assigned to the nearest “synapse”, *i.e*., to the midpoint between Syp and Homer spots, within cut-off distance of 1.5 μm. Such assignments were then categorized as presynaptic or postsynaptic, if they were inside of 45° cones emanating from the midpoint towards Syp and Homer spots, respectively. N = 5 devices per condition from two independent experiments. Paired t-test; * *p* < 5×10^-4^; \*\**p* < 5×10^-5^.

**Figure S11.**
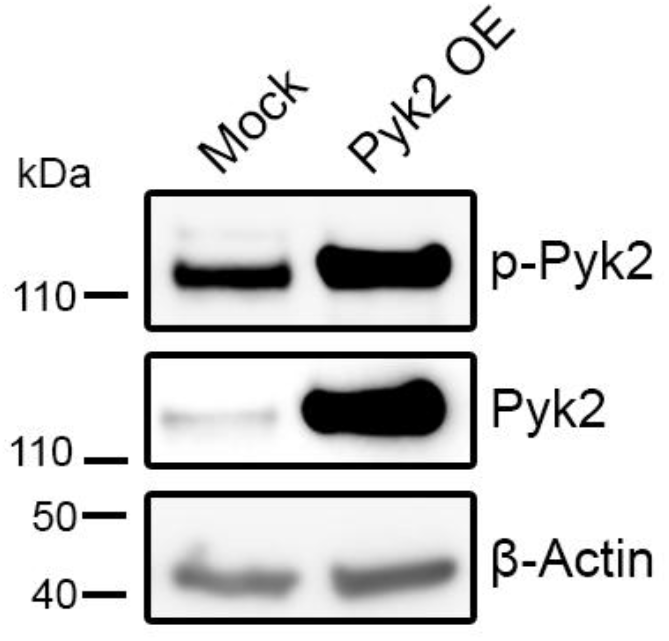
An exemplary immunoblot demonstrating the overexpression of Pyk2 in primary neuronal cultures at DIV14 following lentiviral transduction at DIV7.

**Figure S12.**
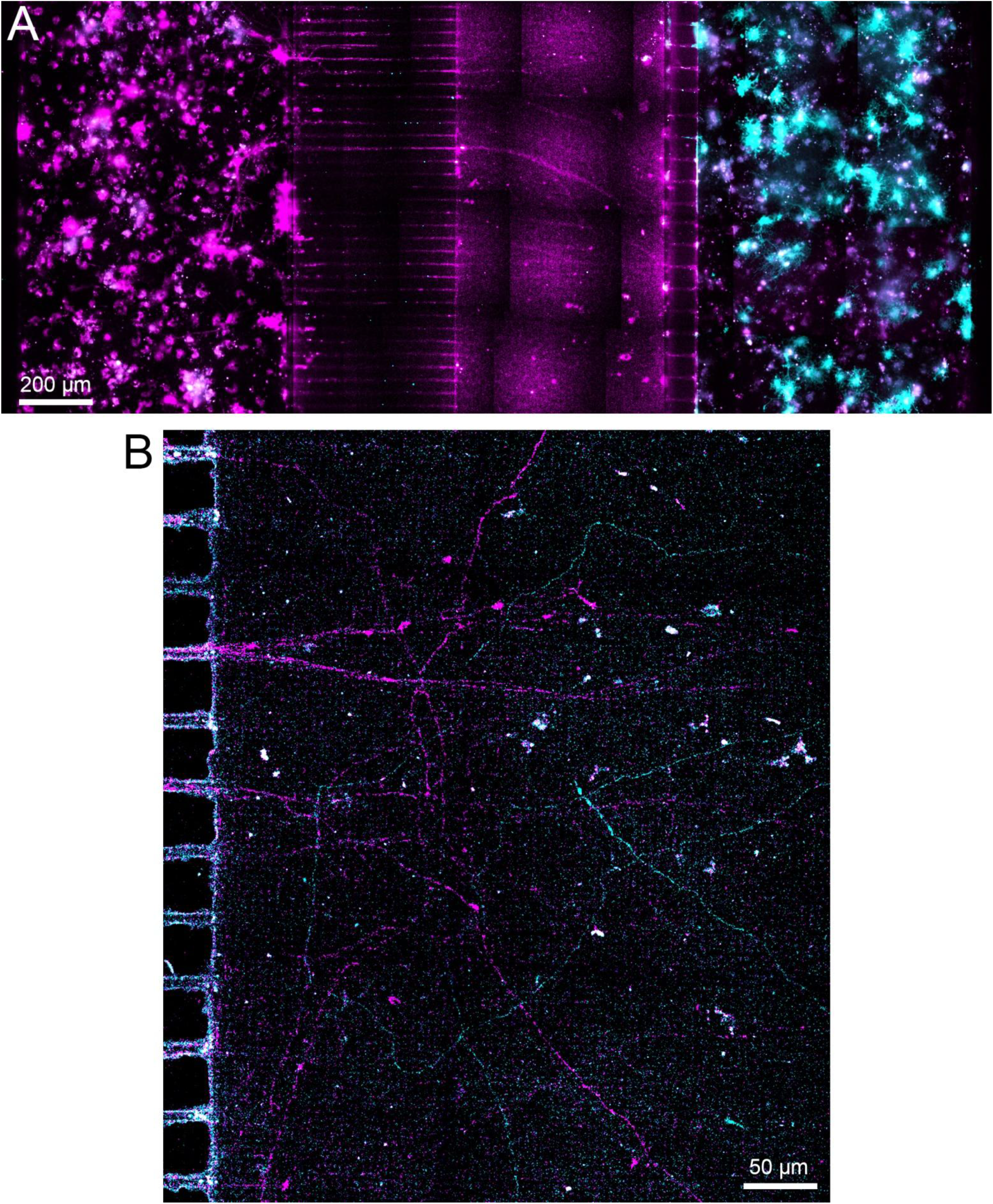
Demonstration of isolated lentiviral treatments at DIV14. **A.** Live-cell microscopy of primary neuron cultures plated in pre- and postsynaptic chambers and transducted at DIV7 with lentiviruses to express LifeAct-ruby (magenta) and LifeAct-GFP (cyan), respectively. **B.** An exemplary area from the synapse chamber, showing fluorescence signals in neurites emanating from both chambers.

## Notes

### Competing Interest Statement

The authors have declared no competing interest.

### Summary of Updates

Additional data and analysis are provided.

